# Highly efficient generation of canker resistant sweet orange enabled by an improved CRISPR/Cas9 system

**DOI:** 10.1101/2021.10.20.465103

**Authors:** Xiaoen Huang, Nian Wang

## Abstract

Sweet orange (*Citrus sinensis*) is the most economically important species for the citrus industry. However, it is susceptible to many diseases including citrus bacterial canker caused by *Xanthomonas citri* subsp. citri (*Xcc*) that triggers devastating effects on citrus production. Conventional breeding has not met the challenge to improve disease resistance of sweet orange due to the long juvenility and other limitations. CRISPR-mediated genome editing has shown promising potentials for genetic improvements of plants. Generation of biallelic/homozygous mutants remains difficult for sweet orange due to low transformation rate, existence of heterozygous alleles for target genes and low biallelic editing efficacy using the CRISPR technology. Here, we report improvements in the CRISPR/Cas9 system for citrus gene editing. Based on the improvements we made previously (dicot codon optimized Cas9, tRNA for multiplexing, a modified sgRNA scaffold with high efficiency, CsU6 to drive sgRNA expression), we further improved our CRISPR/Cas9 system by choosing superior promoters (CmYLCV or CsUbi promoter) to drive Cas9 and optimizing culture temperature. This system was able to generate a biallelic mutation rate of up to 89% for Carrizo citrange and 79% for Hamlin sweet orange. Consequently, this system was used to generate canker resistant Hamlin sweet orange by mutating the effector binding element (EBE) of canker susceptibility gene *CsLOB1*, which is required for causing canker symptoms by *Xcc*. Six biallelic Hamlin sweet orange mutant lines in the EBE were generated. The biallelic mutants are resistant to *Xcc*. Biallelic mutation of the EBE region abolishes the induction of *CsLOB1* by *Xcc*. This study represents a significant improvement in sweet orange gene editing efficacy and generating disease resistant varieties via CRISPR-mediated genome editing. This improvement in citrus genome editing makes genetic studies and manipulations of sweet orange more feasible.

## Introduction

Citrus is one of the most important fruit crops worldwide because of its delightful flavor and scent, as well as health properties. Sweet orange (*Citrus sinensis*) is the most economically important species for the citrus industry. However, citrus production is facing many biotic (e.g., diseases, and insects), and abiotic challenges (e.g., drought, flooding, acidity, salinity, heat, cold, drought, and nutrient deficits) (Li, 2009). Among them, citrus diseases such as citrus canker caused by *Xanthomonas citri* subsp. citri (*Xcc*) (Ference *et al.*, 2018) and Huanglongbing (HLB, also known as citrus greening) caused by *Candidatus* Liberibacter asiaticus (Bové, 2006, Wang, 2019, Yuan *et al.*, 2020) are causing devastating effects on the citrus industry. Most elite citrus varieties, including sweet orange and grapefruit varieties, are susceptible to canker and HLB disease. Genetic improvements of citrus via conventional approaches are challenging. Most citrus varieties result from human selection of natural mutations or natural hybridization rather than artificial hybridization. Hybridization based breeding for citrus is hindered by long juvenile stage (3-10 years), a highly heterozygous nature, heterozygous nature of nucellar seedlings (clones of the female parent), and male or female sterility (Omura and Shimada 2016). It has been suggested that CRISPR-mediated genome editing will revolutionize the genetic improvements of citrus and other tree crops (Dutt *et al.*, 2020, Wheatley and Yang, 2020).

Citrus genome editing mediated by the CRISPR technology was first reported in 2014 (Jia and Wang, 2014a, Jia and Wang, 2014b). The first biallelic mutant of citrus (Carrizo citrange, a rootstock variety) was reported in 2017 (Zhang *et al.*, 2017a) The first biallelic/homozygous mutations of citrus scion varieties (Pummelo (*Citrus maxima*)) were reported in 2020 (Jia and Wang, 2020). It is noteworthy that Pummelo is highly homozygous (Wu *et al.*, 2018) and relatively easy to generate biallelic/homozygous mutants compared to other citrus species. However, most elite citrus varieties including sweet orange are highly heterozygous and biallelic/homozygous mutants have not been reported to date or showed very low editing efficiency (Jia *et al.*, 2017a, Peng *et al.*, 2017, Jia *et al.*, 2019a). We aimed to further improve the CRISPR/Cas system to make genome editing of elite citrus varieties more achievable and improve the disease resistance of sweet orange against canker. Utilization of resistant varieties is the most efficient and eco-friendly approach to control diseases. However, most commercial citrus varieties including sweet orange are susceptible to citrus canker (Favaro *et al.*, 2020, Ference *et al.*, 2020).

Improvement of genome editing efficiency of plants has been conducted via multiple approaches including improvements of the expression of Cas proteins and sgRNA using different promoters. Several promoters for driving Cas9 expression have been tested, such as 35S promoter, ubiquitin promoter (Castel *et al.*, 2019), cell division-specific Yao promoter (Yan *et al.*, 2015), egg cell-specific EC1.2/DD45 (Wang *et al.*, 2015) and RPS5a promoter (Tsutsui and Higashiyama, 2017b, Ordon *et al.*, 2020). Among them, the ubiquitin promoter, and RPS5a promoter have significantly improved genome editing efficacy compared to the commonly used 35S promoter in Arabidopsis, tomato, rice or maize (Stavolone *et al.*, 2003, Cermak *et al.*, 2017, Tsutsui and Higashiyama, 2017a, Castel *et al.*, 2019). Endogenous polymerase III promoters (U3 or U6) improve the expression of sgRNAs (Sun *et al.*, 2015, Qi *et al.*, 2018). In addition, other approaches including codon optimization of the Cas proteins (Li *et al.*, 2014), and temperature optimization (Xiang *et al.*, 2017) have shown promises to improve the efficacy of genome editing.

*Xcc* causes the characteristic hypertrophy and hyperplasia symptoms on citrus tissues via secretion of PthA4, a transcriptional activator-like (TAL) effector, through the type III secretion system (Swarup *et al.*, 1991, Yan and Wang, 2012, An *et al.*, 2020). PthA4 enters the nucleus and activates the expression of the canker susceptibility gene *LATERAL ORGAN BOUNDARIES 1* (*LOB1*) via binding to the effector binding element (EBE) in the promoter region using its central nearly identical tandem repeats of 33-34 amino acids (Hu *et al.*, 2014). The specific recognition between nucleotides and the repeat is determined by the 12^th^ and 13^th^ amino acids of each repeat, known as the “repeat-variable diresidue” (RVD) (Moscou and Bogdanove, 2009, Römer *et al.*, 2009). Mutation of the EBE regions has been used to generate disease resistance crops against TAL effector-dependent pathogens such as bacterial blight of rice (Li *et al.*, 2012, Oliva *et al.*, 2019). In our previous study, genome modified Duncan grapefruit of the coding region of *LOB1* demonstrates canker resistance (Jia *et al.*, 2017b). Mutation of both alleles of the EBE region of *LOB1* is required to abolish its induction by PthA4 and generate canker resistant citrus plants (Jia *et al.*, 2016, Jia and Wang, 2020, Jia *et al.*, 2021). However, biallelic/homozygous mutation of the EBE region for elite varieties such as sweet orange is yet to be achieved. Most importantly, highly efficient generation of biallelic/homozygous mutation in elite varieties such as sweet orange and grapefruit has not been reported yet.

In this study, we improved CRISPR/Cas9 system, which was used to generate six biallelic *CsLOB1* EBE mutant lines of Hamlin sweet orange. All biallelic *CsLOB1* EBE mutant plants (100%) are resistant against *Xcc*, representing a milestone in canker resistance development for elite citrus varieties. In addition, this improved CRISPR/Cas9 system efficiently (100%) edits the tobacco genome. Hence, this improved CRISPR/Cas9 system may be applied to other dicots besides citrus and tobacco.

## Results

### Improvement of CRISPR/Cas9 system for citrus genome editing

First, we sought to improve the Cas9/CRISPR-mediated genome editing efficiency in citrus using the phytoene desaturase (*PDS*) gene as a target. The *PDS* gene was chosen for editing because of the obvious albino phenotype caused by homozygous or biallelic knockout mutation of the *PDS* gene (Zhang *et al.*, 2017a, Huang *et al.*, 2020). The albino phenotype serves as the functional editing (null mutation) readout. We first edited Carrizo citrange, a hybrid between *Citrus sinensis* ‘Washington’ sweet orange and *Poncirus trifoliata*, which can be relatively easily transformed (compared to sweet orange) and edited based on our previous experience (Jia *et al.*, 2017a, Huang *et al.*, 2020). A GFP visual maker was included in the constructs to further confirm successful plant transformation in addition to antibiotic marker selection. Inspired by previous reports showing that increased culture temperature can increase editing efficacy (LeBlanc *et al.*, 2018, Malzahn *et al.*, 2019, Milner *et al.*, 2020), we sought to test if we can increase editing efficacy by increasing culture temperature. The editing efficiency was higher at 30°C for both gene editing in protoplasts and in *Agrobacterium*-mediated transformed epicotyl tissue than that at room temperature (Supplementary Fig. 1).

Previously, we improved the CRISPR/Cas9 system by codon optimizing the Cas9 gene, identifying citrus U6 promoter (CsU6), using an improved gRNA scaffold and inclusion of tRNA for multiplexing (Xie *et al.*, 2015, Huang *et al.*, 2020). Here we aimed to select a superior promoter for driving *Cas9* expression. The 320 bp core region of the 35S promoter in the CRISPR/Cas9 construct was replaced with either the *Citrus sinensis* ubiquitin (CsUbi) or CmYLCV promoter (Supplementary Data 1). The CsUbi promoter (Cs4g11190) was selected through mining of citrus RNAseq data (Ribeiro *et al.*, 2021). Cs4g11190 is highly expressed in various stages of leaf development (Supplementary Table 1). The CmYLCV promoter (Stavolone *et al.*, 2003) is a strong promoter and functions in both monocots and dicots. The upstream 538 bp regulatory region containing the enhancer elements of 35S was kept and fused with candidate promoters (Fig.1A) to enhance Cas9 expression and/or promote early expression of Cas9 in first dividing cells (Ow *et al.*, 1987, Benfey *et al.*, 1990). The editing efficacy of 35S, CsUbi, and CmYLCV driving Cas9 expression was compared (Fig. 1A). The albino phenotype was used as a readout for null mutations (Fig. 1B). The 35S promoter produced 50% albino plants; the CsUbi promoter and CmYLCV promoter generated 80% and 89% albino plants, respectively (Fig. 1C, Supplementary Fig. 2), suggesting that CsUbi and CmYLCV promoters are superior to 35S for gene editing in citrus.

**Fig. 1.**
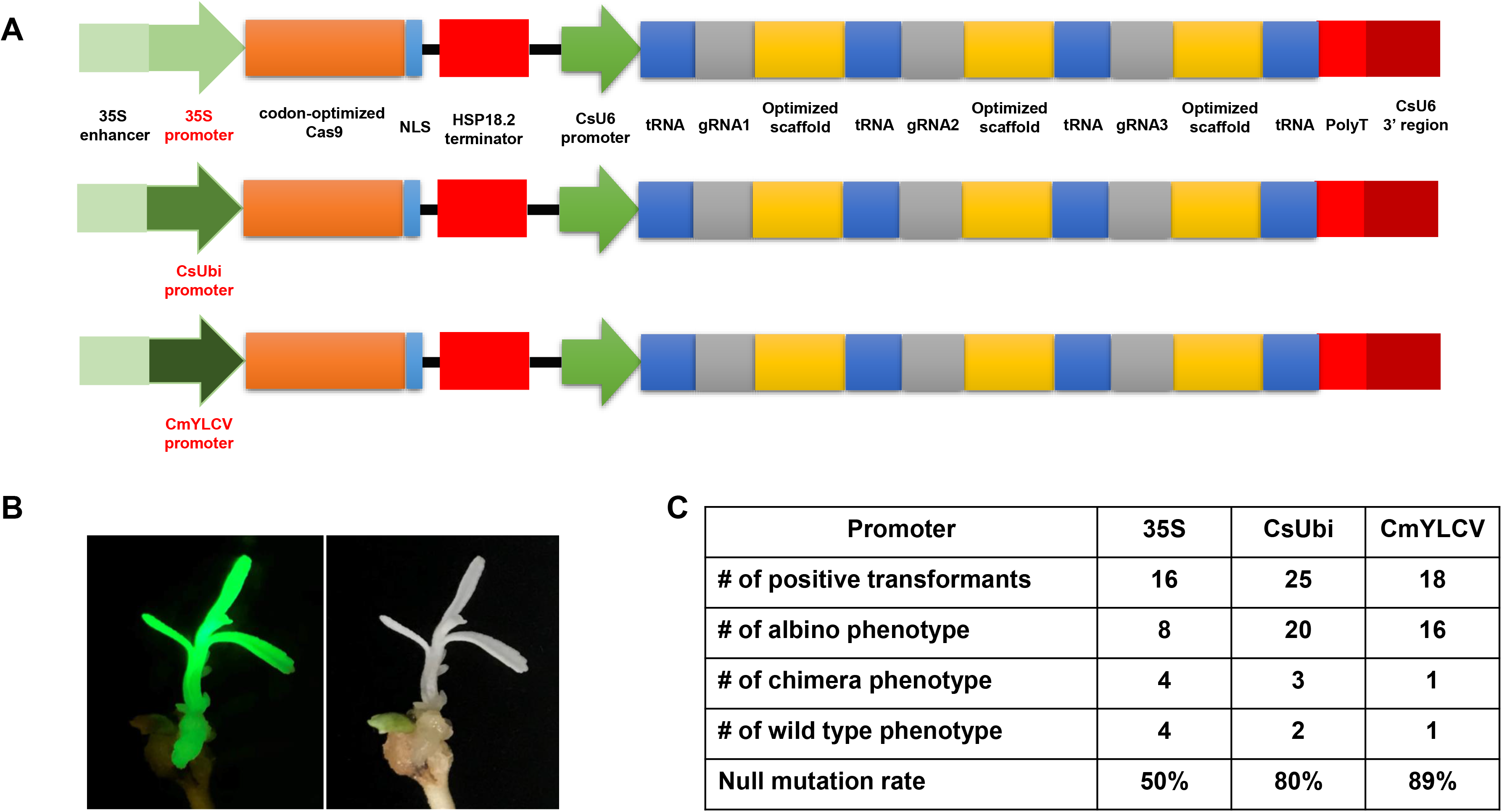
Optimal promoters for driving Cas9 expression to improve gene editing efficacy in citrus. **A)** 35S promoter, citrus Ubiquitin promoter (CsUbi) and *Cestrum yellow leaf curling virus* (CmYLCV) promoter for driving Cas9 expression in the CRISPR constructs. **B)** A representative picture showing albino phenotype, caused by null mutation of the *PDS* gene with CmYLCV promoter construct. The construct contains GFP as visual marker. **C)** Summary of editing efficacy from three constructs shown in A.

In all transformation events, we observed chimeric albino phenotype. It is common to observe chimera phenotypes during plant transformation of explants, such as epicotyledons, leaves, and stems. The chimera rate for the 35S, CsUbi and CmYLCV was 25%, 12%, 5.6%, respectively (Figure 1C). These results suggest that CsUbi and CmYLCV promoters drive an early expression of the Cas9 in the first transformed cells compared to 35S promoter. All constructs contained GFP. Not surprisingly, GFP florescence was observed to colocalize with albino phenotype. Therefore, for all the transformation experiments, only transformants with evenly distributed GFP florescence all over the plantlets were kept for further analyses.

### The improved CRISPR/Cas9 system efficiently edits tobacco (*Nicotiana tabacum*)

The CmYLCV promoter is a strong constitutive promoter for heterologous gene expression in *Arabidopsis thaliana* and *Nicotiana tabacum* and in a wide variety of crops including *Lycopersicon esculentum*, *Zea mays* and *Oryza sativa* (Stavolone *et al.*, 2003). We hypothesized that the improved CRISPR/Cas9 system driven by CmYLCV works in other plant species, especially dicots. To test this hypothesis, we tested the construct in tobacco (*N. tabacum*), an allotetraploid species, considering its amenability to stable transformation. There are two *PDS* homologs, *PDS-A* and *PDS-B* in tobacco genome. Two gRNAs were designed in the conserved regions between the two homologs (Fig. 2A). We obtained a total of 11 transformants with green fluorescence (GFP). The transformants with green fluorescence all showed albino phenotype (Fig. 2B, Supplementary Fig. 3), indicating that we achieved tetra-allelic mutations for *PDS-A* and *PDS-B* in all positive transformants (Fig.2C). This result demonstrates that the improved CRISPR/Cas9 system achieves a 100% null mutation efficacy for *PDS* genes in tobacco. This result suggests that our improved CRISPR/Cas9 system may be able to efficiently edit genomes of other dicots, besides citrus and tobacco.

**Fig. 2.**
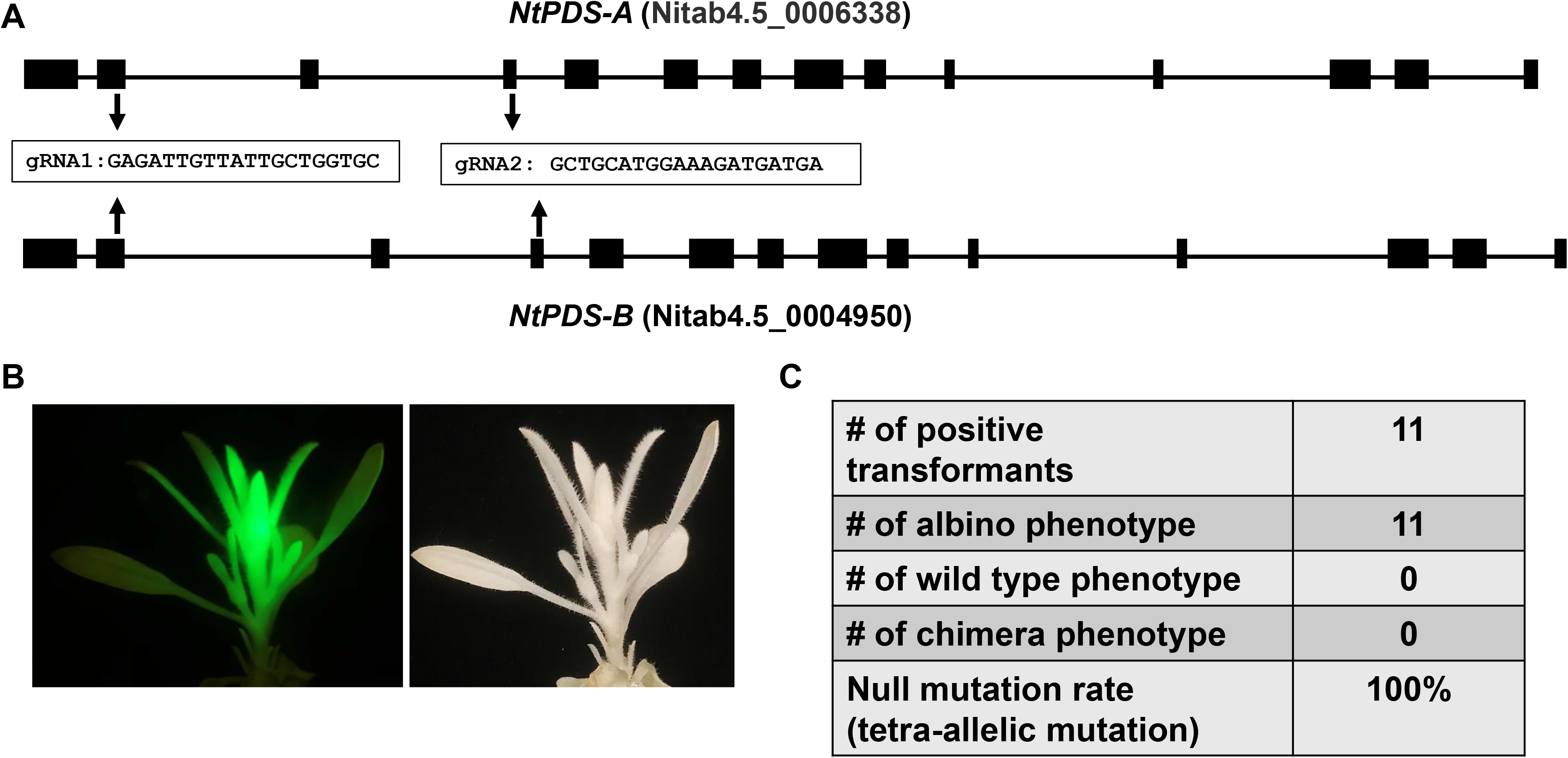
The improved CRISPR/Cas9 system works efficiently for genome editing of tobacco (*Nicotiana tabacum*). **A)** Gene structures of two *NtPDS* homologs from allotetraploid *Nicotiana tabacum*. Two gRNAs conserved in both *NtPDS-A* and *NtPDS-B* genes as indicated were designed for genome editing. **B)** A representative picture shows the albino phenotype in genome modified *Nicotiana tabacum*. The construct contains GFP as visual marker. **C)** Summary of editing efficacy.

### Highly efficient generation of biallelic mutation in sweet orange

Sweet orange is a hybrid between pummelo (*Citrus maxima*) and mandarin (*Citrus reticulata*) and most genes have two different alleles. Here, we used the improved CRISPR/Cas9 system that contains a codon optimized Cas9, tRNA for multiplexing (Xie *et al.*, 2015, Huang *et al.*, 2020), CsU6 promoter (Huang *et al.*, 2020), improved sgRNA scaffold (Dang *et al.*, 2015) and the CmYLCV promoter to edit the EBE in the promoter region of the canker susceptibility gene *CsLOB1*(Hu *et al.*, 2014). The EBE region of sweet orange contains two different types/alleles. We aimed to edit both alleles to create null mutations.

Two different gRNAs for both type I and type II EBEs were designed. A third gRNA targeting a region upstream of the translation start codon ATG was also included (Fig. 3A). The final construct contains these three gRNAs. Such a design aimed to create not only small indels, but also longer deletion mutations. We first transformed citrus protoplasts to confirm the functionality of the construct. As expected, the construct not only produced indel mutations, but also generated relatively long deletion mutations (Fig. 3B, C, D).

**Fig. 3.**
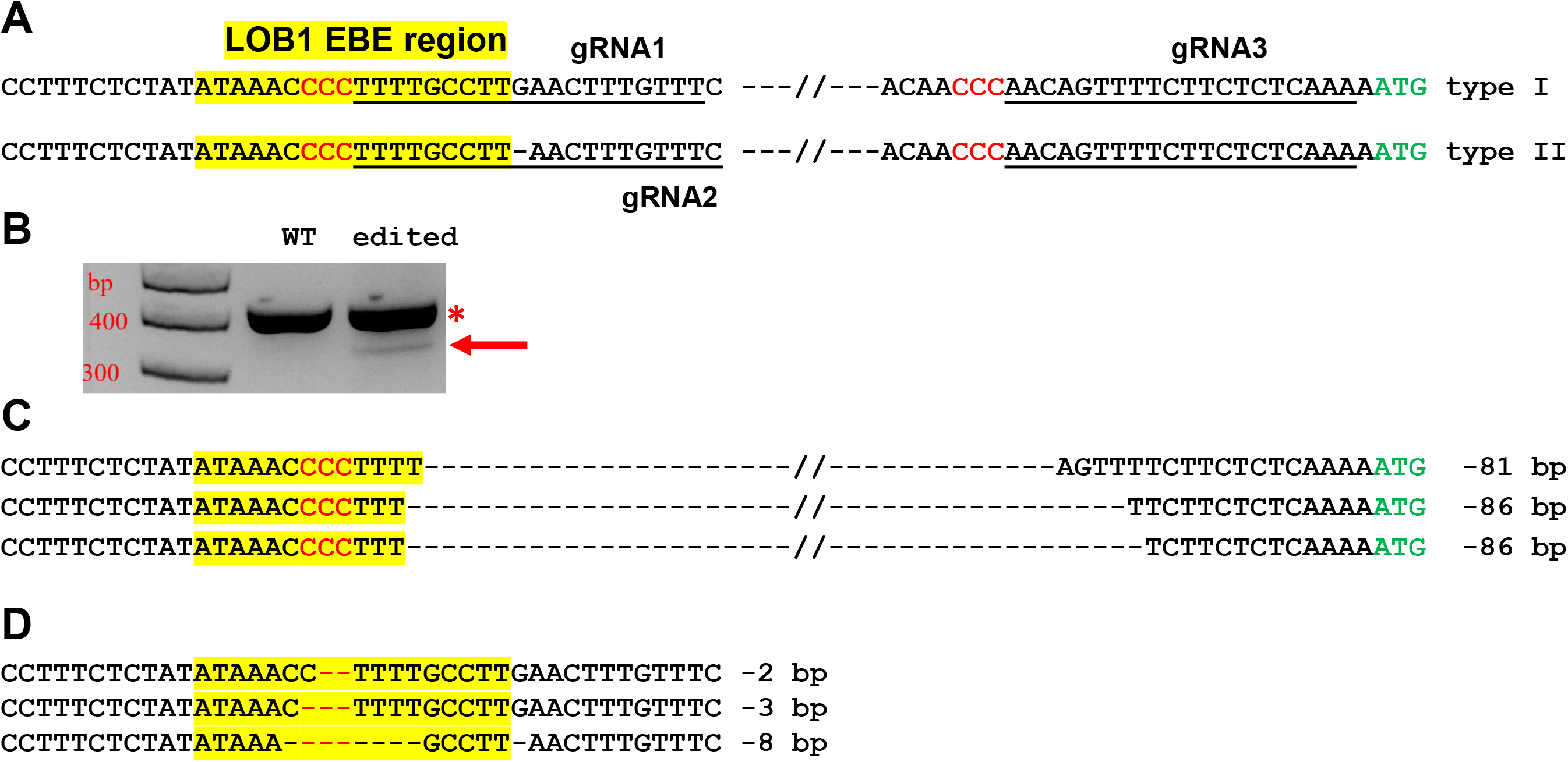
*CsLOB1* editing in Hamlin sweet orange protoplasts. **A)** *CsLOB1* promoter contains two different types. The EBE region was highlighted with yellow; regions chosen for gRNAs were underlined; PAM was shown in red. The final CRISPR constructs contains 3 gRNAs as indicated. **B)** PCR products generated by two primers spanning the EBE and last gRNA. The PCR product with an arrow was amplified from the deletion mutants. **C)** Sanger sequencing results of the PCR product with the arrow in C. **D)** Sanger sequencing results of the PCR products marked by * in B that were cloned for colony sequencing.

Next, we performed stable transformation of Hamlin sweet orange epicotyls. A total of 16 positive transformants were obtained. Two lines died during regeneration. Finally, we obtained 14 transgenic plants. We genotyped these 14 seedlings. Among them, 11 plants contained biallelic mutations in the EBE region (Fig. 4, Supplementary Fig. 4), representing a 78.6% biallelic mutation rate (Supplementary Table 2). The 3 heterozygous mutants contained both wild type and edited EBEs. Subsequently, we micro-grafted these seedlings onto the rootstock variety Carrizo citrange. A total of 6 biallelic mutants and 2 heterozygous mutants were successfully grafted on the rootstock, whereas the rest did not survive.

**Fig. 4.**
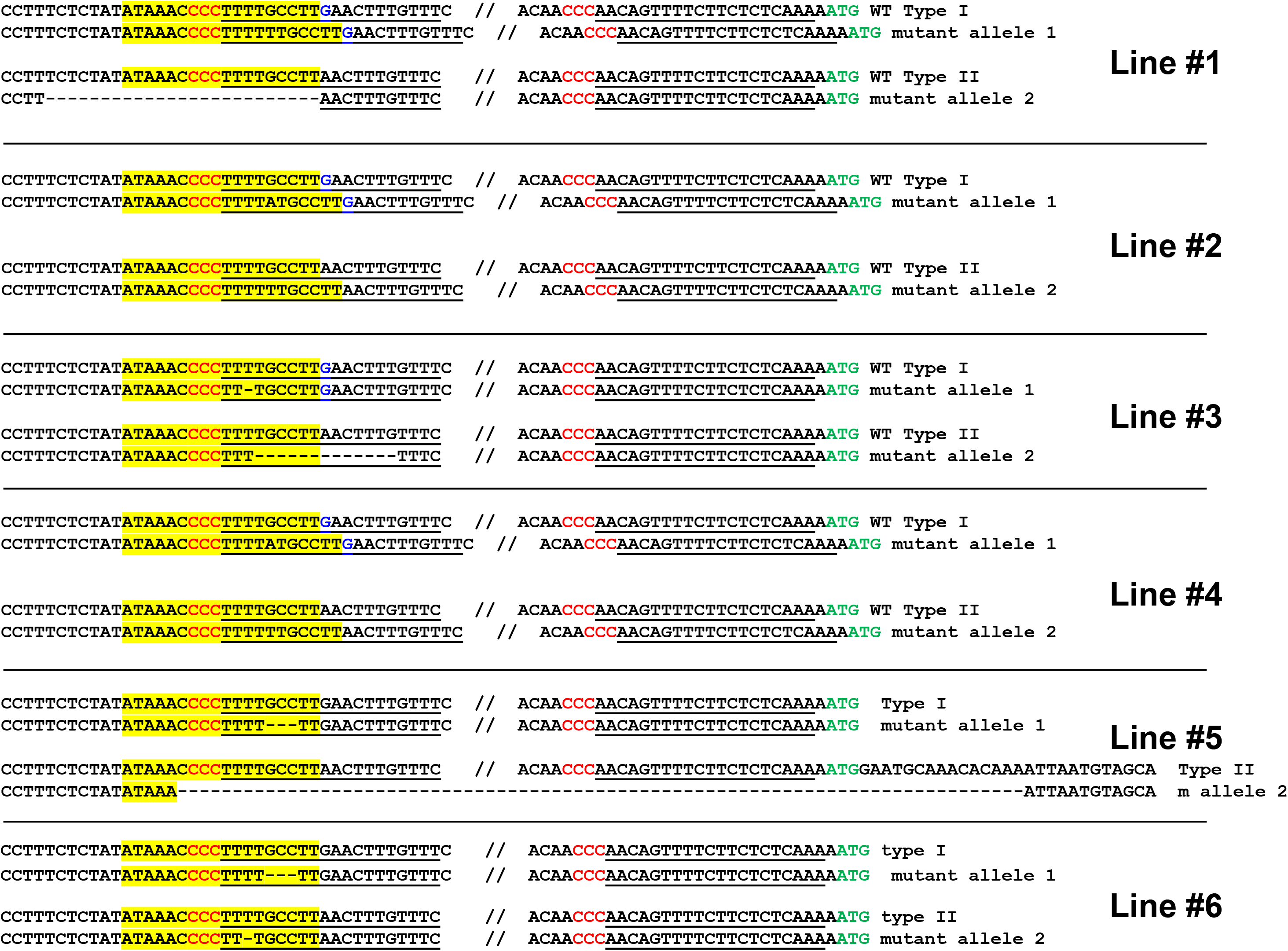
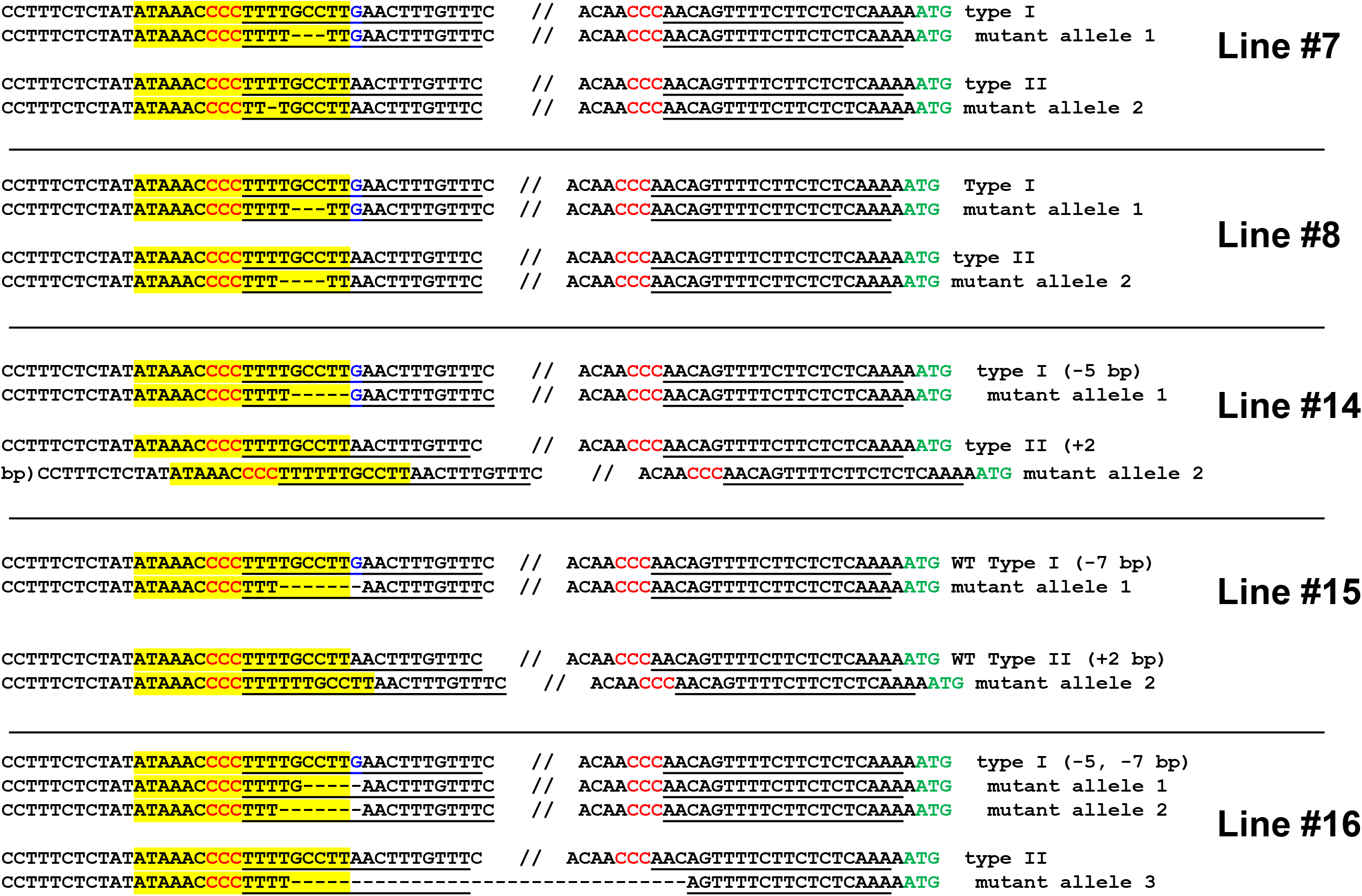
Highly efficient generation of biallelic mutant citrus plants for Hamlin sweet orange. Genotyping of biallelic mutants. Mutations of the EBE region is the focus of the genotyping. The EBE region was highlighted with yellow; regions chosen for gRNAs were underlined; PAM was indicated in red; nucleotide G in blue is a SNP only in one allele, but not in another allele.

### Biallelic mutants are resistant to *Xcc*

We hypothesized that the biallelic mutation of the EBE region of *CsLOB1* disrupts its binding to *Xcc* TAL effector PthA4, thus *Xcc* is unable to induce canker susceptibility gene *CsLOB1* for symptom development (Swarup *et al.*, 1992, Hu *et al.*, 2014). To test this hypothesis, wild type Hamlin sweet orange plants, one heterozygous mutant line (line #11) and six biallelic mutant lines (line #1, 2, 3, 4, 5, 15) of Hamlin sweet orange were tested for resistance against *Xcc* strains (Fig. 5). For each genotype, one half of the leaf was inoculated with wild type *Xcc*, and another half was inoculated with *Xcc pthA4*:Tn5 mutant carrying the designer TALE dLOB2, which activates the expression of *LOB2* (Teper *et al.*, 2020). *LOB2* is a *LOB1* homolog that causes canker symptoms when artificially induced, such as in the presence of dLOB2 (Zhang *et al.*, 2017b, Teper *et al.*, 2020). *Xcc pthA4*:Tn5 dLOB2 was included as a control. The onset of canker symptoms by *Xcc pthA4*:Tn5 dLOB2 on the same leave can exclude the possibility that lack of canker symptoms for biallelic mutants is owing to leaf age. At 8 dpi, all wild type Hamlin leaves developed canker lesions for both wild type *Xcc* and dLOB2 carrying *Xcc pthA4*:Tn5. No canker symptoms were observed for the biallelic mutant lines (line #1, 2, 3, 4, 5 and 15) when inoculated with wild type *Xcc*. On the contrary, canker symptoms were observed on the other half of the leaves inoculated with *XccpthA4:Tn5* dLOB2. The heterozygous mutant line #11 showed canker disease symptoms for both wild type *Xcc* and dLOB2 carrying *Xcc pthA4*:Tn5 (Fig. 5). Taken together, the present results demonstrate that the biallelic Hamlin sweet orange mutant lines are resistant to *Xcc*.

**Fig. 5.**
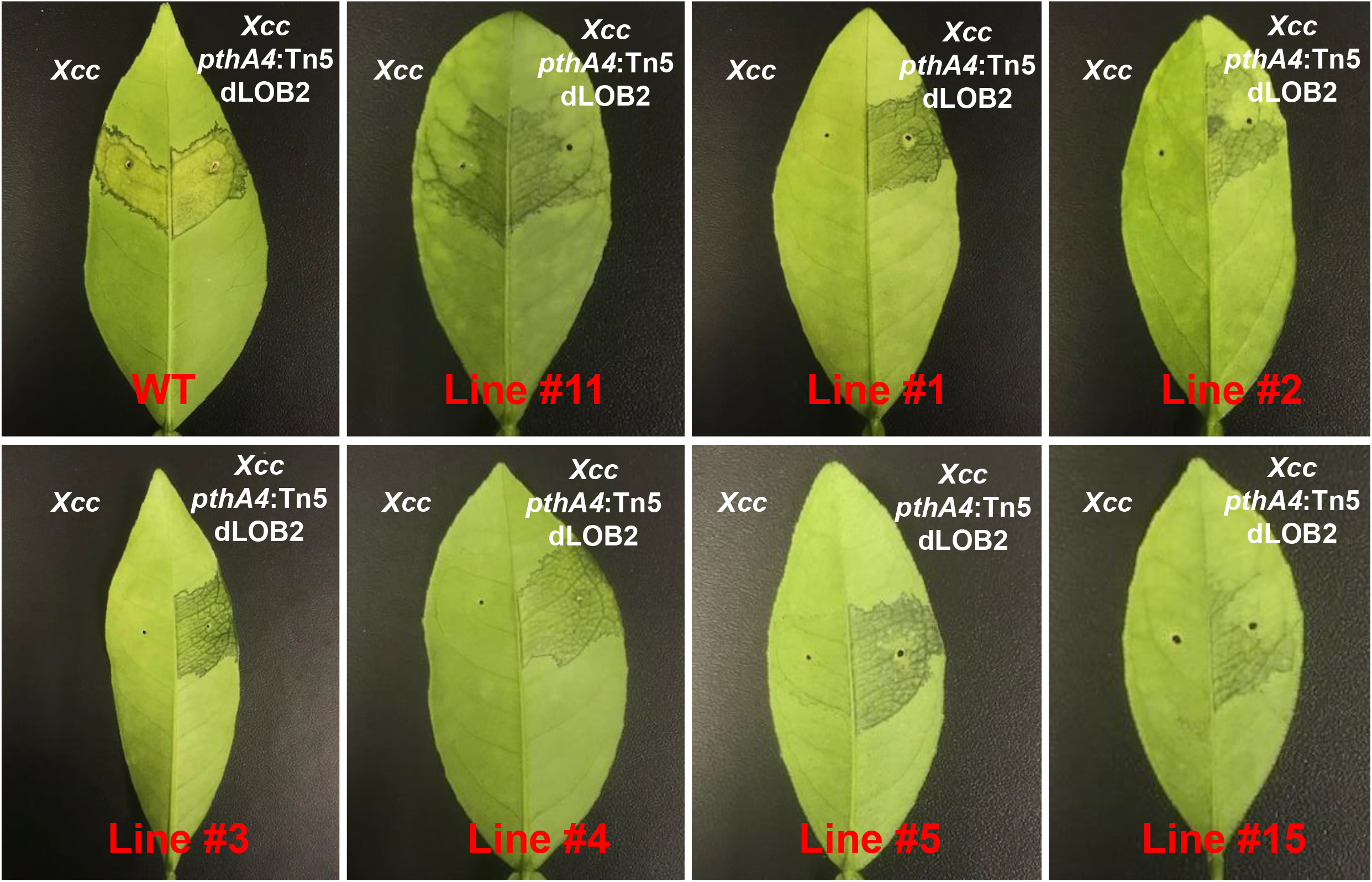
Biallelic mutants of the EBE region of *CsLOB1* of Hamlin sweet orange are resistant to *Xcc.* *Xcc* inoculation assay was conducted using fully expanded young leaves. Line #11 is a heterozygous mutant; other lines are biallelic mutants. For each line, left half of the tested leaf was syringe inoculated with wild type *Xcc* (10^8^ CFU/ml); the right half of the tested leaf was inoculated with *Xcc pthA4:Tn5* carrying designer TAL effector dLOB2 (10^8^ CFU/ml). Pictures were taken at 8 days post inoculation.

### *Xcc* does not induce the expression of *CsLOB1* in biallelic mutants

Next, we tested whether biallelic mutations of the EBE region of *CsLOB1* abolish its induction by *Xcc* using reverse transcription-quantitative PCR (RT-qPCR). The expression level of *CsLOB1* in wild type Hamlin sweet orange was dramatically induced when inoculated with *Xcc*, which is consistent with previous findings (Hu *et al.*, 2014, Teper *et al.*, 2020). The induction of *CsLOB1* was obliterated in the biallelic mutant line Ham1 (Fig. 6A). *Xcc pthA4:Tn5 dLOB2* induced the expression of *LOB2* (Fig. 6B and C), which is consistent with canker symptoms induced by *Xcc pthA4:Tn5 dLOB2* on both wild type and biallelic mutant lines (Fig. 5) and with our previous result (Teper *et al.*, 2020).

**Fig. 6.**
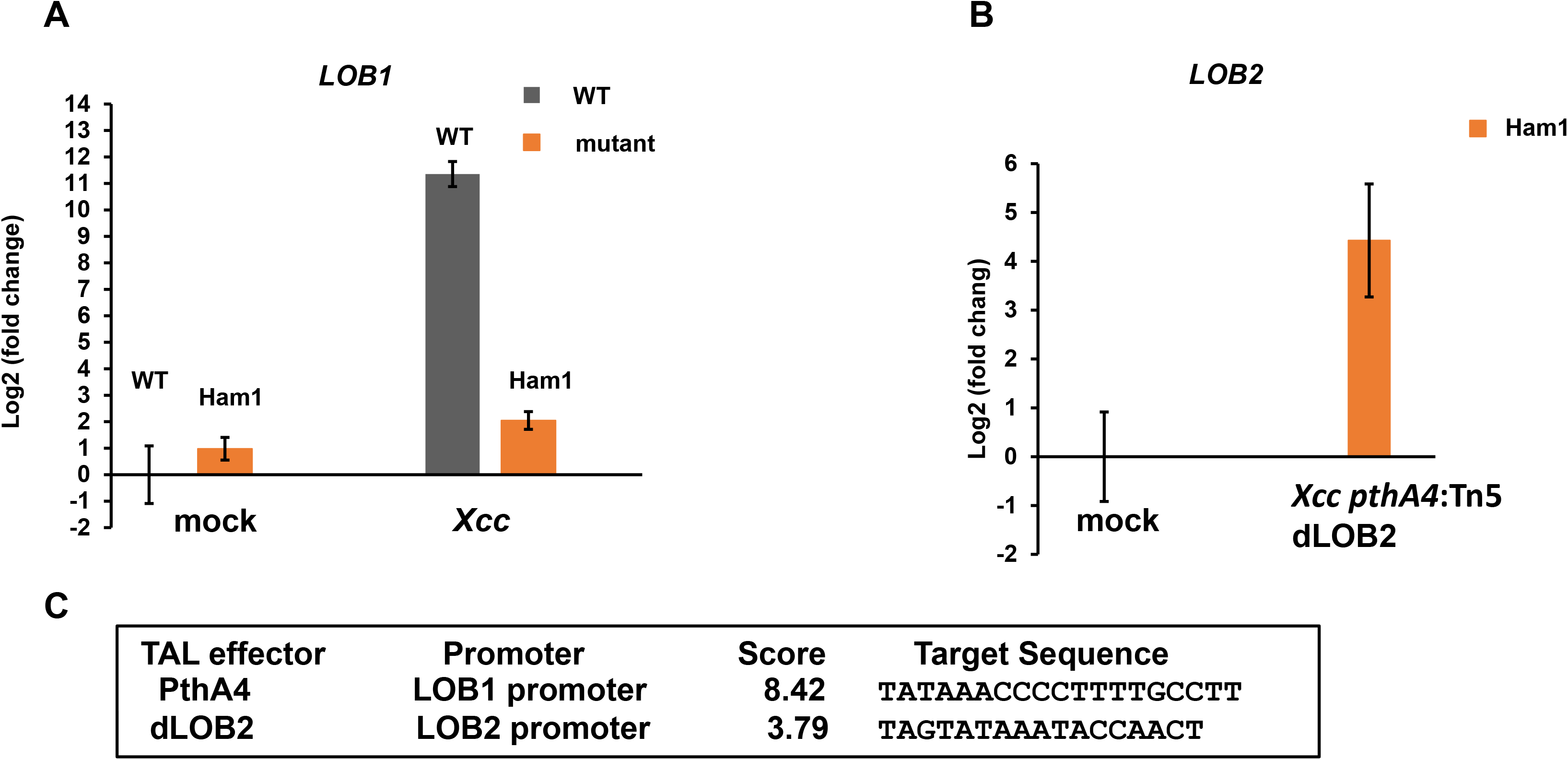
Induction of *CsLOB1* by *Xcc* is abolished in a biallelic mutant line of Hamlin sweet orange. **A)** RT-qPCR analyses of Cs*LOB1* relative expression in leaf samples collected at 48 hours post inoculation with indicated treatments. Mock, mock inoculation with buffer; *Xcc*, inoculation with wild type *Xcc* (10^8^ CFU/mL); dLOB2, inoculation with *Xcc pthA4:Tn5* carrying designer TAL effector dLOB2 (10^8^ CFU/mL). Each treatment has three biological repeats. Ham1, a representative biallelic mutant line #1 (aka Ham1). All expression levels were normalized to the WT mock. The GAPDH gene was used as an endogenous control. **B)** RT-qPCR analysis of *LOB2* relative expression of leaf samples collected at 48 hours post inoculation with indicated treatment. All expression levels were normalized to the Ham1 mock. The GAPDH gene was used as an endogenous control. **C)** Finding promoter binding site for TAL effectors by using Target Finder (https://tale-nt.cac.cornell.edu/node/add/talef-off). Designer TAL effector dLOB2 shows affinity to the LOB2 promoter, but not the LOB1 promoter. “/” indicates no affinity.

## Discussion

In this study we improved CRISPR/Cas9 system to achieve high efficacy for biallelic editing in citrus. The improved system contains dicot codon optimized Cas9, tRNA for multiplexing, an improved sgRNA scaffold with high efficiency (Dang *et al.*, 2015), CsU6 to drive sgRNA expression, and superior promoters (CmYLCV or CsUbi promoter). Importantly, this improved CRISPR/Cas9 system has significantly increased the biallelic/homozygous mutation rates. For Carrizo citrange, the CsUbi and CmYLCV driven constructs achieved null mutation rates of 80 and 89%, respectively, via epicotyl transformation. In addition, the CmYLCV construct attained 79% biallelic mutation rate in Hamlin sweet orange even under restrained gRNA options (EBE region). Choices of optimal gRNAs through online tools, such as CRISPR-P and CRISPR-PLANT, may further boost biallelic mutation rate (Liu *et al.*, 2017, Minkenberg *et al.*, 2019). The difference in editing efficacy may also results from the differential transformation efficacy in different citrus genotypes. The transformation efficacy was reported to be 20.6% for Carrizo citrange(Pena *et al.*, 1995), but ranged from 0.88 to 23.8 for sweet orange varieties (Poles *et al.*, 2020) and was 2.5% for Hamlin sweet orange (Fávero *et al.*, 2012). Previously, the biallelic mutation rate was reported to be 5.3% for grapefruit (Jia *et al.* 2021), 44.4% for Carrizo citrange (11.1% homozygous mutation rate) (Huang *et al.*, 2020) and 22.7% to 55% for Carrizo citrange (Zhang *et al.*, 2017a). Thus, this improved CRISPR/Cas9 system we reported here represents a significant improvement compared to previous studies. In addition, this new system uses a citrus U6 promoter and a CmYLCV promoter to drive sgRNA or Cas9 protein, respectively. It has been recommended to use endogenous promoters to drive Cas proteins or sgRNA to improve efficacy or to address gene regulation concerns. It was reported that U6 promoters and two ubiquitin promoters identified in grapevine significantly increase the editing efficiency in grape by improving the expression of sgRNA and Cas9, respectively (Ren *et al.*, 2021). Similarly, endogenous promoters for U6 and ubiquitin were successfully used for genome editing of maize and yielded mutation efficiencies ranging from 48.5% to 97.1% (Qi *et al.*, 2018). The improved CRISPR/Cas9 system developed in this study can be used not only for editing the genome of citrus, but also for other plant species. We successfully edited tobacco *PDS* genes (100% null mutation) with the CmYLCV-driven construct. We anticipate that the modified CRISPR/Cas9 system we developed is efficient in genome editing of other plant species, especially dicots.

The CmYLCV promoter and CsUbi promoter driving the expression of Cas9 have significantly improved the citrus editing efficacy compared to that of the 35S promoter. The CmYLCV promoter was reported to function in both dicots and monocots and shows higher or comparable activities than the 35S promoter in driving the heterozygous expression (Stavolone *et al.*, 2003). Similar as the 35S promoter, the CmYLCV promoter is an RNA polymerase II promoter. The CmYLCV promoter demonstrates similar efficacy as the Ubiquitin 2 promoter of switchgrass in transformation, reaching about 80% and 100% of protoplasts after 24-hours and 48-hours, respectively, post transformation (Weiss 2020). In addition to driving the expression of Cas proteins, the CmYLCV promoter has also been successfully used to drive the sgRNA array via Csy4-mediated processing or gRNA-tRNA array (Čermák 2017).

We successfully obtained multiple biallelic canker resistant Hamlin sweet orange mutant lines against *Xcc*, which is a TAL effector-dependent pathogen, via editing the EBE region of canker susceptibility gene *LOB1* (Hu *et al.*, 2014). TAL effectors are critical virulence factors for most *Xanthomonas* pathogens (Boch and Bonas, 2010, Perez-Quintero and Szurek, 2019, An *et al.*, 2020). TAL effectors are known to be subject to repeated recombination that allows them to recognize new targets or overcome the resistance resulting from mismatches between the TAL effectors and the EBE regions. It was reported that *Xcc* is able to overcome 2-5 mismatches between the repeat region of PthA4 and the EBE of *CsLOB1*, but is unable to overcome the resistance when the mismatch is equal or more than 7 (Teper and Wang, 2021). However, the previous study was conducted via an accelerated evolutionary approach that includes injection with high titers of *Xcc* than normally encountered in natural settings. In addition, the TALEs were cloned into pBBR1MCS5, a medium copy number vector (approximately 30 copies), whereas naturally occurring PthA4 is three copies in each bacterial cell (Teper and Wang, 2021). It is anticipated that the chance for *Xcc* to overcome the mismatch between PthA4 and the EBE region in natural settings is much lower than that observed via the accelerated evolutionary study (Teper and Wang, 2021). Additionally, the genome modified lines will enable us to investigate whether *Xcc* can overcome the resistance resulting from EBE mutations in natural settings in future studies. The biallelic mutants also can serve as a diagnostic tool to monitor emerging *Xcc* strains that target other susceptibility genes such as *LOB2* and *LOB3* (Zhang *et al.*, 2017b, Teper *et al.*, 2020). A similar strategy has been developed for *Xoo*/rice pathosystem (Eom *et al.*, 2019).

Of note, the canker resistant sweet orange lines contain the CRISPR/Cas construct, which is considered as transgenic and might not be suitable to be used for citrus production. Nevertheless, successful generation of canker resistant sweet orange represents a milestone in the ultimate utilization of canker resistant citrus varieties based on mutations of the coding or the EBE region of the canker susceptibility gene *LOB1*. Generation of canker resistant citrus varieties will overcome the significant issues in canker management using copper-based products (Zhang *et al.*, 2003), antibiotics (Hu and Wang, 2016, Li *et al.*, 2019, Li *et al.*, 2020), and biocontrol bacteria. Such an approach has been successfully used to generate disease resistant rice varieties (Oliva *et al.*, 2019). For rice, the CRISPR/Cas construct can be easily removed by progeny segregation or back-crossing, which is not practical for citrus. This is because citrus has long juvenile stage. In addition, back-crossing leads to the loss of the elite quality of parental varieties. In future studies, we will aim to generate non-transgenic genome modified canker resistant citrus varieties via transient expression of the CRISPR/Cas construct. Such a method has been successfully used to generate nontransgenic genome-edited crops (Zhang *et al.*, 2016, Chen *et al.*, 2018, Lin *et al.*, 2018, Veillet *et al.*, 2019). Development of methods using in vitro assembled CRISPR/Cas9/gRNA ribonuclearprotein (Cas9/gRNA RNP) complexes for transient expression of the CRISPR/Cas9 system in citrus may also be an attractive future option given that this approach has proven successful in producing other crop plants that contain precisely edited genes and that are non-transgenic (Woo *et al.*, 2015, Svitashev *et al.*, 2016, Zhang *et al.*, 2021). In addition, newly emerging DNA delivery technologies such as nanoparticles (Demirer *et al.*, 2019, Lv *et al.*, 2020) might help deliver our highly efficient constructs to achieve nontransgenic genome editing.

## Methods

### Plant materials

The seeds of Carrizo citrange were purchased from Lyn Citrus Seeds (Arvin, California, USA). The seeds of sweet orange Hamlin were harvested from mature fruits in the groves of the Citrus Research and Education Center, University of Florida.

### Constructs

To test performance of different promoters for driving Cas9, we replaced the 320 bp core region of the 35S promoter in construct pC-CsU6-tRNA-3PDS, which was reported previously (Huang *et al.*, 2020), with promoters used in this study. Specifically, the 35S promoter sequence was digested with BspEI and SalI from pC-CsU6-tRNA-3PDS. CsUbi or CmYLCV (Supplementary Data 1) was PCR amplified and inserted into the same site to replace the 35S promoter. PC-CsUbi-PDS and PC-CmYLCV-PDS constructs were obtained and confirmed by Sanger sequencing. To make the *CsLOB1* EBE CRISPR construct, primers LOBPro3-F1/R1 and LOBPro3-F2/R2 (Supplementary Table 3) were used to make multiplex constructs using the transient vector pXH1-CsU6-tRNA via Golden gate cloning as described previously (Huang *et al.*, 2020). Three gRNAs were included: one targeting type I EBE region of *CsLOB1*, one targeting type II EBE region of *CsLOB1* (Jia *et al.*, 2016, Jia *et al.*, 2017b, Jia and Wang, 2020) and one targeting a region upstream of ATG. The construct PXH1-LOB1-Pro3 was confirmed by Sanger sequencing. The CmYLCV promoter sequence (Supplementary Data 1) was cut from PC-CmYLCV-PDS with SphI+SalI and inserted in the same site of PXH1-LOB1-Pro3 to replace the 35S promoter to make the transient expression construct PXH1-CmYLCV-LOB1-Pro3. This construct was used for protoplast transfection.

For stable transformation, the cassette was cut from PXH1-CmYLCV-LOB1-Pro3 with FspI and XbaI and inserted into the ZraI/XbaI site of the binary vector PCXH2, which carries the GFP expression cassette to make the final construct PCXH2-CmYLCV-LOB1-Pro3. This construct, after transforming into *Agrobacterium* strain EHA105 was used for *Agrobacterium*-mediated citrus epicotyl transformation.

For tobacco *NtPDS* gene editing, we also used the same tRNA-based multiplex vector as described above. The Cas9 is driven by CmYLCV promoter, and the gRNAs for *NtPDS* are driven by CsU6 promoter. Two gRNAs were designed (gRNA1: GAGATTGTTATTGCTGGTGC and gRNA2: GCTGCATGGAAAGATGATGA) based on the conserved regions between the two *PDS* homologs in the genome of allotetraploid species *Nicotiana tabacum*. The final CRISPR construct was tr transformed into *Agrobacterium* strain EHA105, which was used for stable transformation of tobacco leaf discs.

### Protoplast transformation

Transformation of CRISPR constructs into embryogenic citrus protoplasts was performed as described previously (Huang *et al.*, 2020).

### Citrus transformation

*Agrobacterium*-mediated transformation of citrus epicotyl was performed by following the protocol described previously (Jia *et al.*, 2019b, Huang *et al.*, 2020). After co-cultivation of epicotyls with *Agrobacterium*, the transformed epicotyls were incubated in regeneration medium at 30°C or at room temperature in the dark for two weeks before culturing under light at room temperature. For Carrizo *PDS* gene editing, two independent experiments were performed.

### Genotyping of citrus transformants

Since the transformation constructs carry the GFP expression cassette, positive transformants were selected under GFP filter in addition to antibiotic selection (Kanamycin). Citrus genomic DNA was extracted using the CTAB (cetyl trimethylammonium bromide) method (Pandey and Wang, 2019). Positive transformants were further confirmed by Cas9 amplification (primers in Supplementary Table 3). Detection of editing in the EBE region of *CsLOB1* promoter was performed by amplifying the target region (primers in Supplementary Table 3) with high fidelity DNA polymerase Q5 (New England Biolabs, Ipswich, MA, USA), followed by cloning of PCR products and sequencing.

### *Xcc* inoculation

*Xanthomonas citri* subsp. *citri* (*Xcc*) wild type strain 306 and dLOB2 containing *Xcc pthA4*:Tn5 (Yan and Wang, 2012, Hu *et al.*, 2016, Teper *et al.*, 2020) were suspended in 20 mM MgCl2 at 10^8^ CFU/ml. The bacteria suspensions were syringe-infiltrated into young fully expanded leaves of wild type sweet orange Hamlin sweet orange plants or mutant plants (Teper and Wang, 2021). Inoculated plants were kept in a temperature-controlled (28°C) glasshouse with high humidity. Pictures were taken 8 days post inoculation for disease resistance evaluation.

### RT-qPCR

Leaves of wild type and a biallelic mutant line #1 (hereafter named Ham1) with and without *Xcc* inoculation were sampled at 48 hours post inoculation. Total RNA was extracted using the TRIzol reagent (Invitrogen, Carlsbad, CA, USA) following the manufacturer’s manual. Extracted RNA was further treated with RQ1 RNase-Free DNase (Promega, Madison, WI, USA) to remove genomic DNA contamination, if any. The treated RNA was further purified with RNeasy Mini Kit (QIAGEN, Germantown, MD, USA). One microgram of RNA from each sample was used for cDNA synthesis by using QuantiTect Reverse Transcription Kit (QIAGEN). Quantitative PCR was performed using SYBR Green PCR Master Mix in QuantStudio 3 Real-Time PCR System (Applied Biosystems, Foster City, CA). The citrus house-keeping gene *GAPDH* was used as an endogenous control. Quantification of gene expression was calculated by following the comparative *Ct* method (Pfaffl, 2001). Primers for RT-qPCR are listed in Supplementary Table 2.

## Acknowledgements

We appreciated Dr. Yuanchun Wang for performing micro-grafting. The research has been supported by USDA National Institute of Food and Agriculture grant # 2018-70016-27412, #2016-70016-24833, and #2019-70016-29796, USDA-NIFA Plant Biotic Interactions Program 2017-67013-26527, Florida Citrus Initiative, and Florida Citrus Research and Development Foundation.

## Author contributions

XH and NW designed and organized the study. XH performed the experiments. NW and XH prepared the manuscript.

## Short supporting materials legends

**Supplementary Fig. 1.**
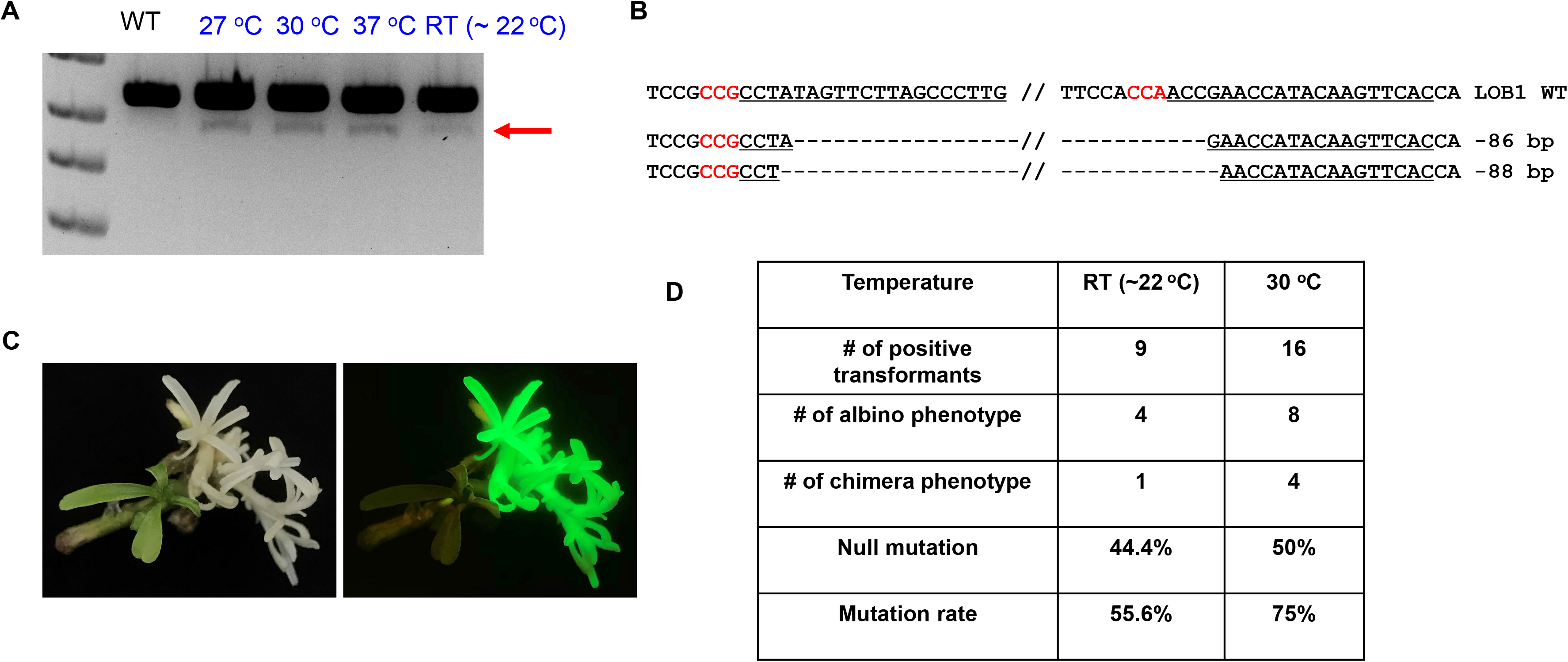
Higher temperatures can increase SpCas9 editing efficacy in citrus. **A)** CRISPR/Cas9-mediated editing of *CsLOB1* in citrus protoplasts under different temperature. Three independent assays were performed, with similar results. **B)** Sanger sequencing result for PCR product indicated with the arrow in A; the sequencing data is from the sample at 30℃. **C)** A representative picture showing albino phenotype. **D)** Summary of editing efficacy at different temperature. Null mutation rate was calculated by dividing the number of pure albino plants with number of total positive transformants. Mutation rate was calculated by dividing the number of pure albino plants and chimeric plants with number of total positive transformants.

**Supplementary Fig. 2.**
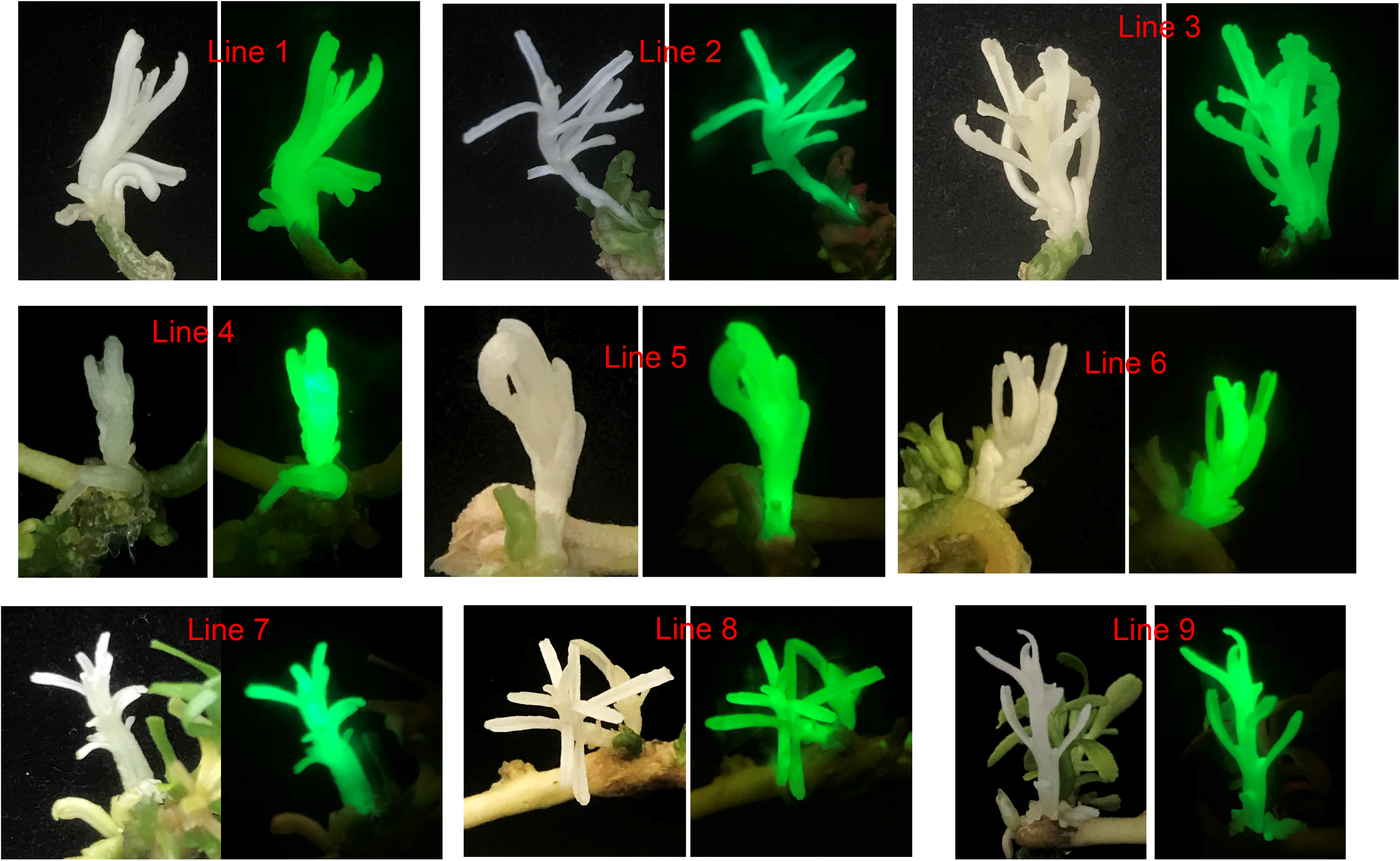
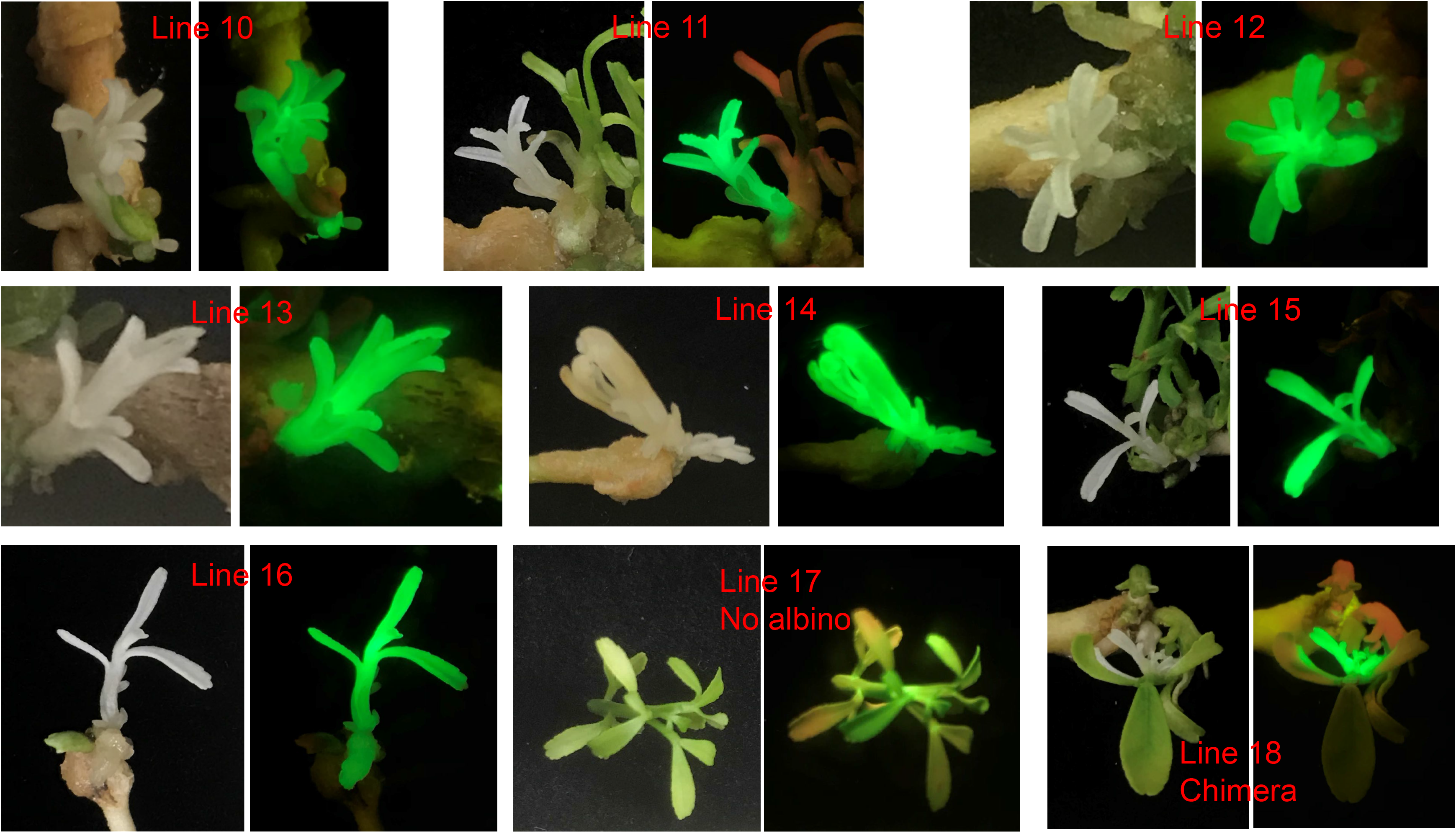
Phenotypes from CmYLCV-Cas9-PDS construct.

**Supplementary Fig. 3.**
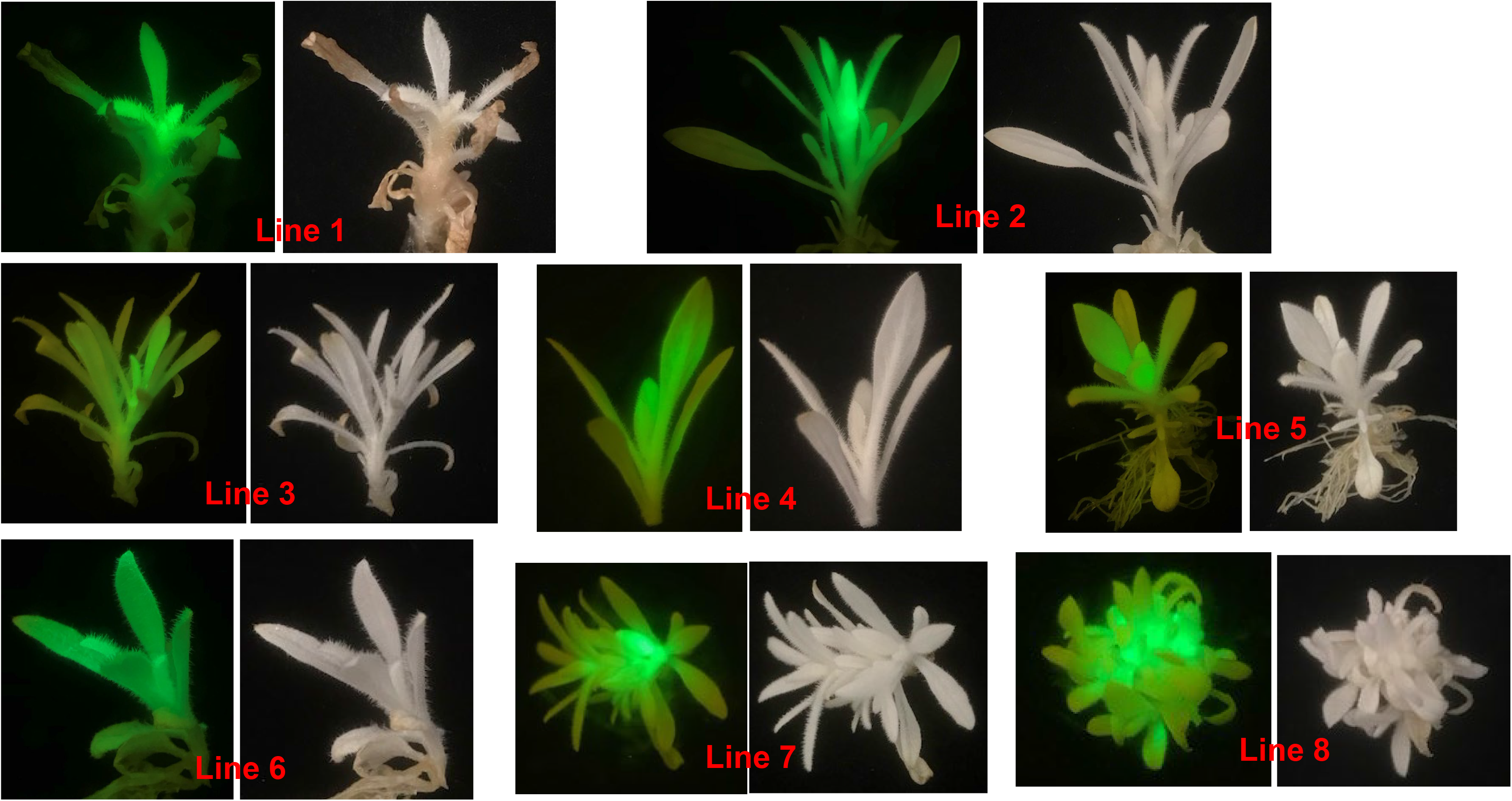
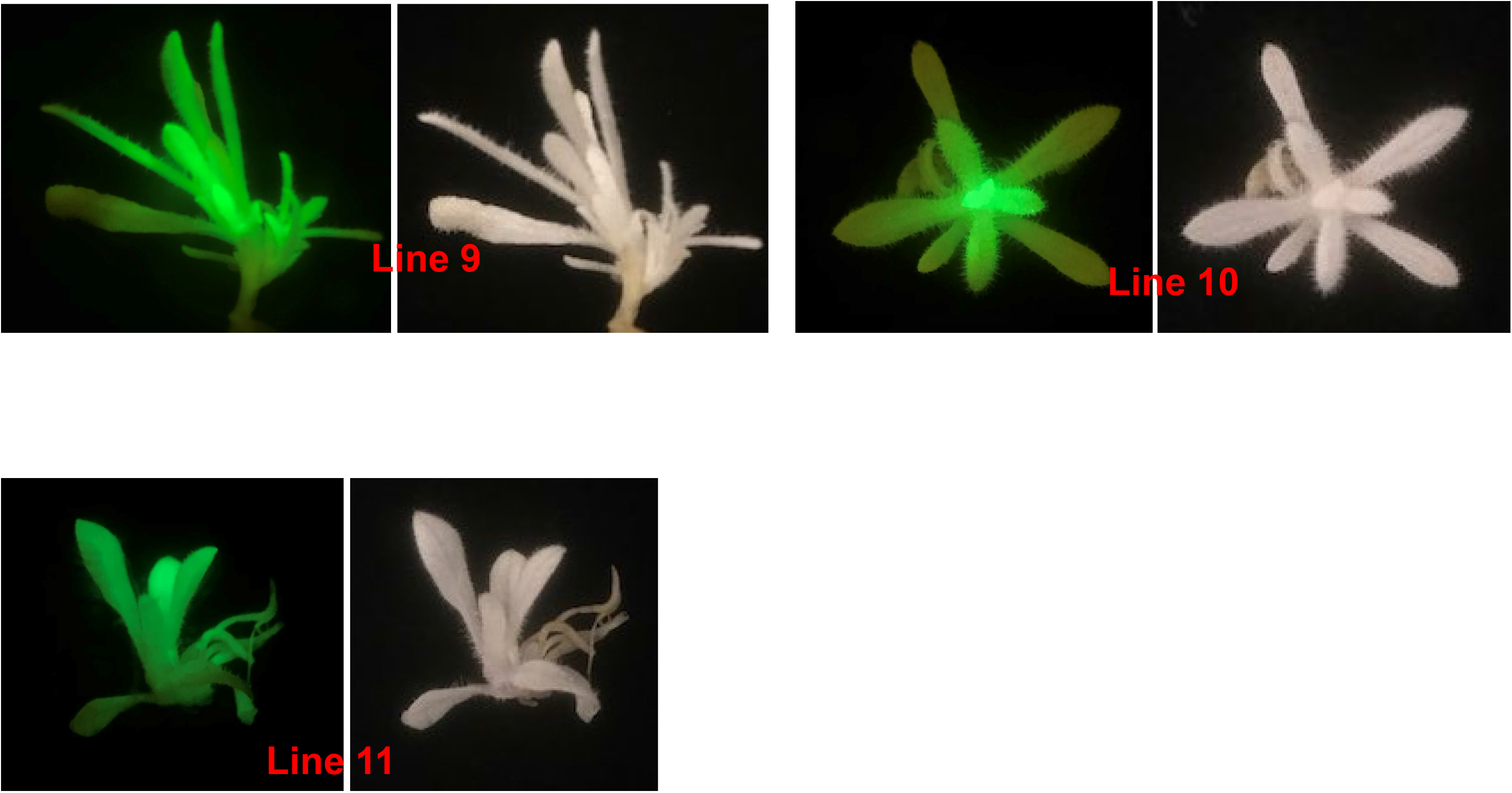
Optimized CRISPR/Cas9 system can efficiently edit *PDS* genes in tobacco (*Nicotiana tabacum*)

**Supplementary Fig. 4.**
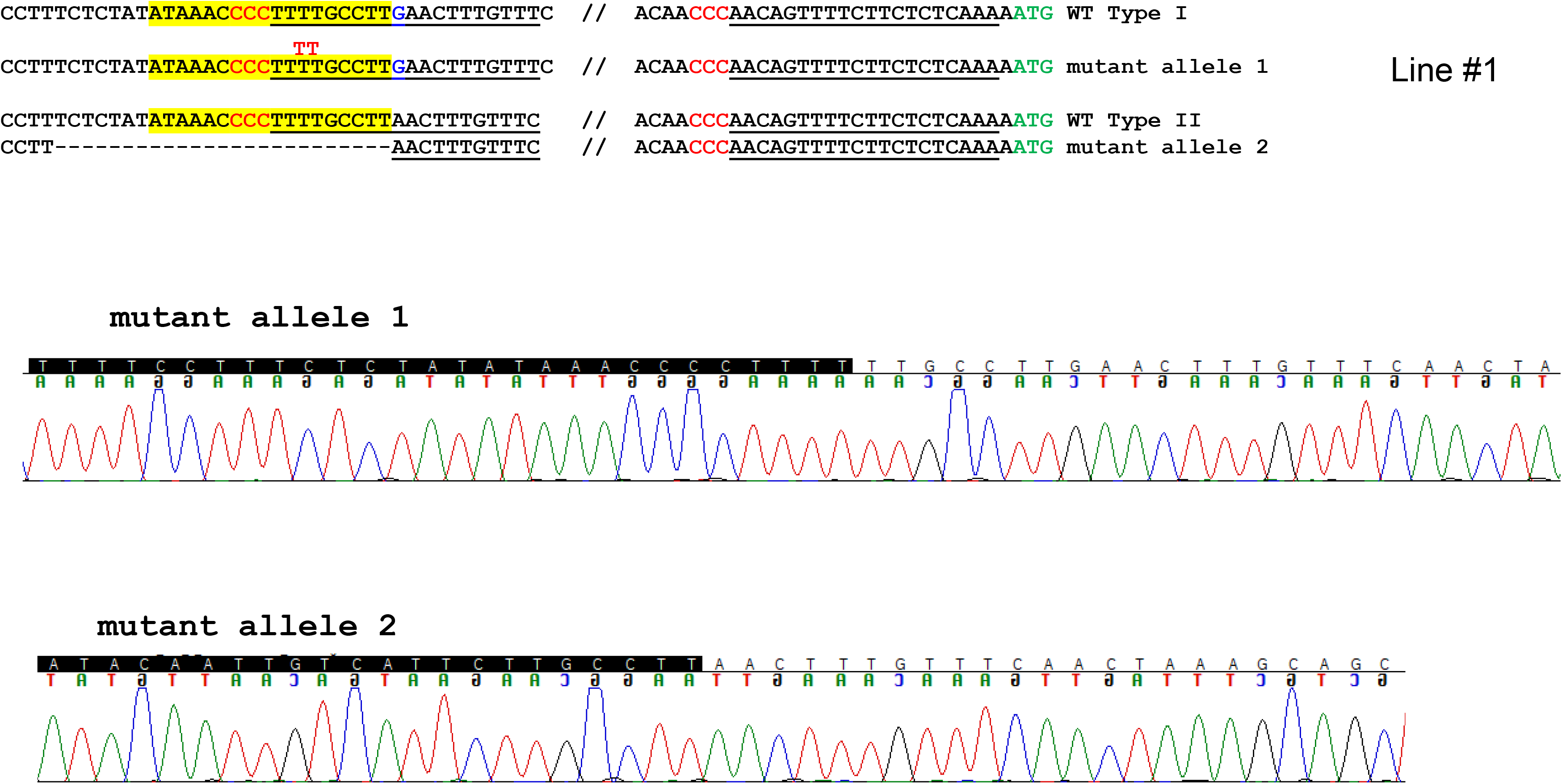
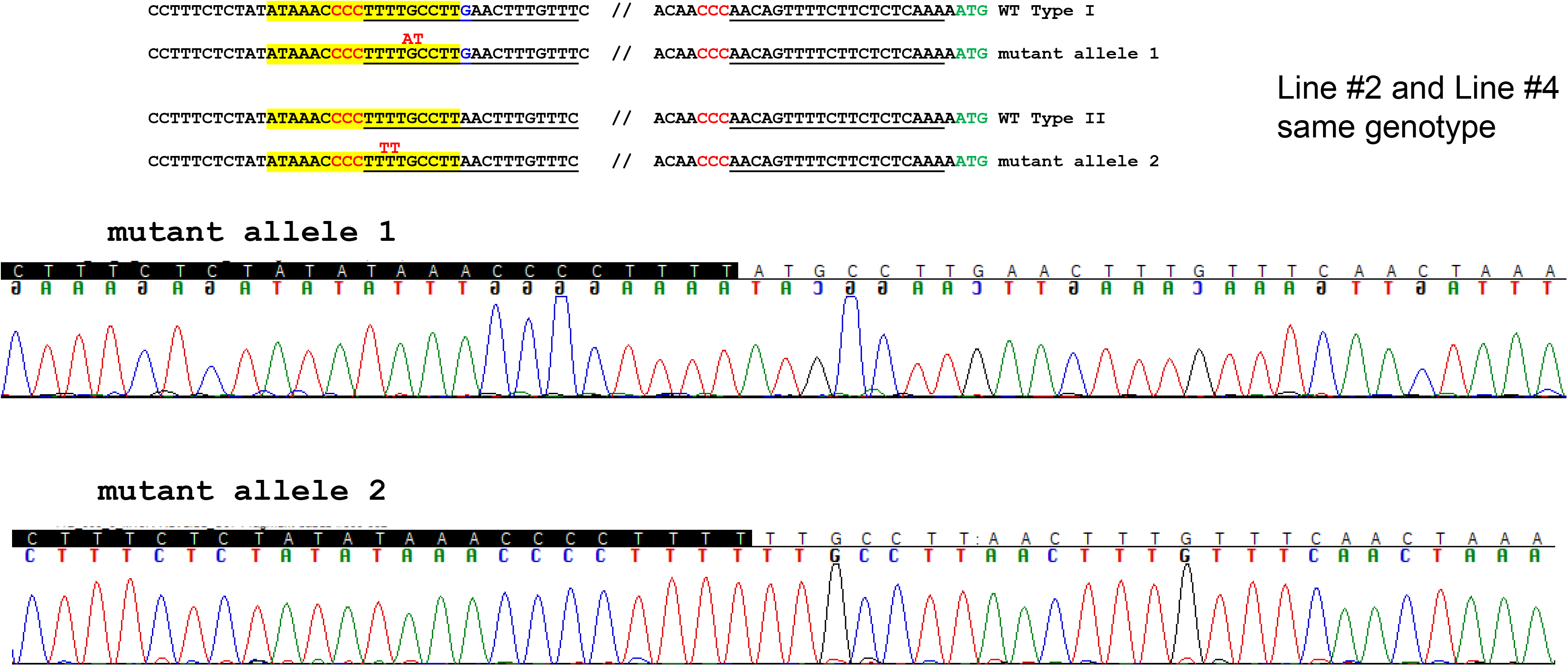
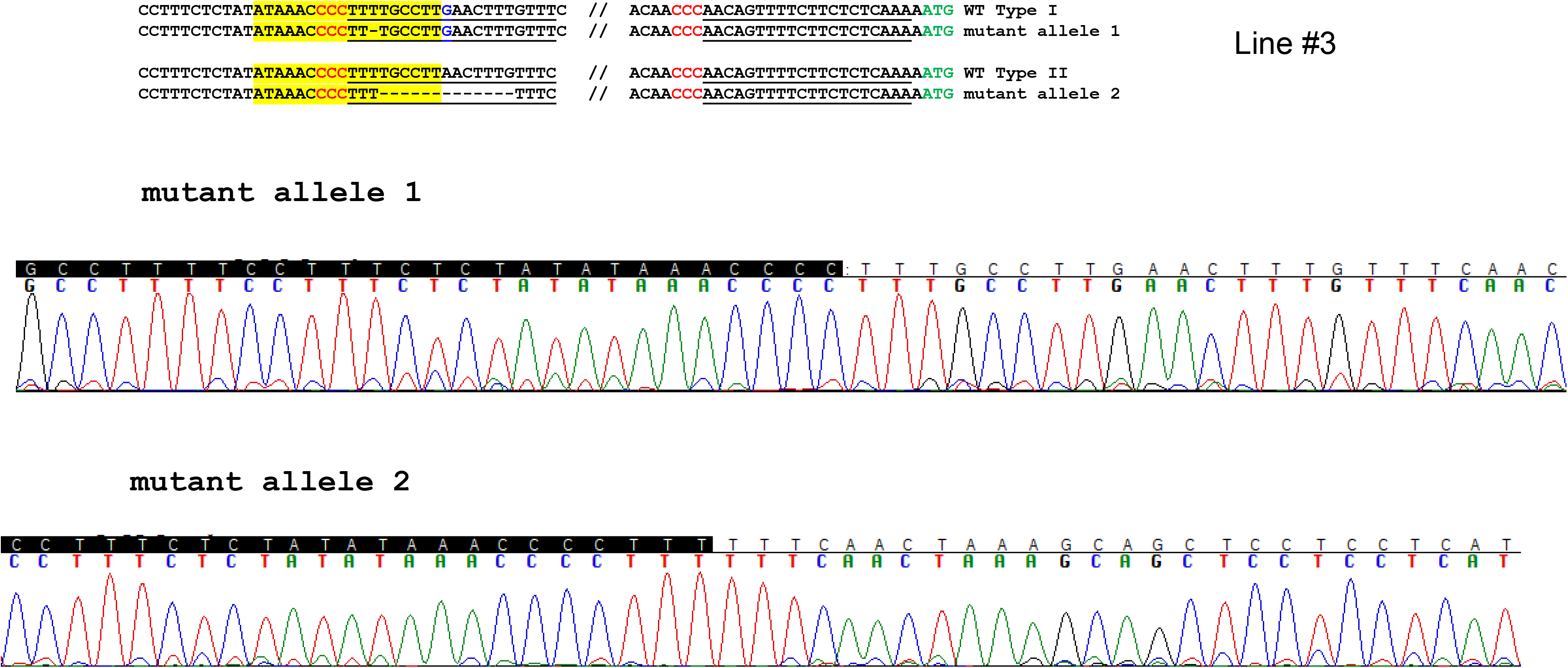
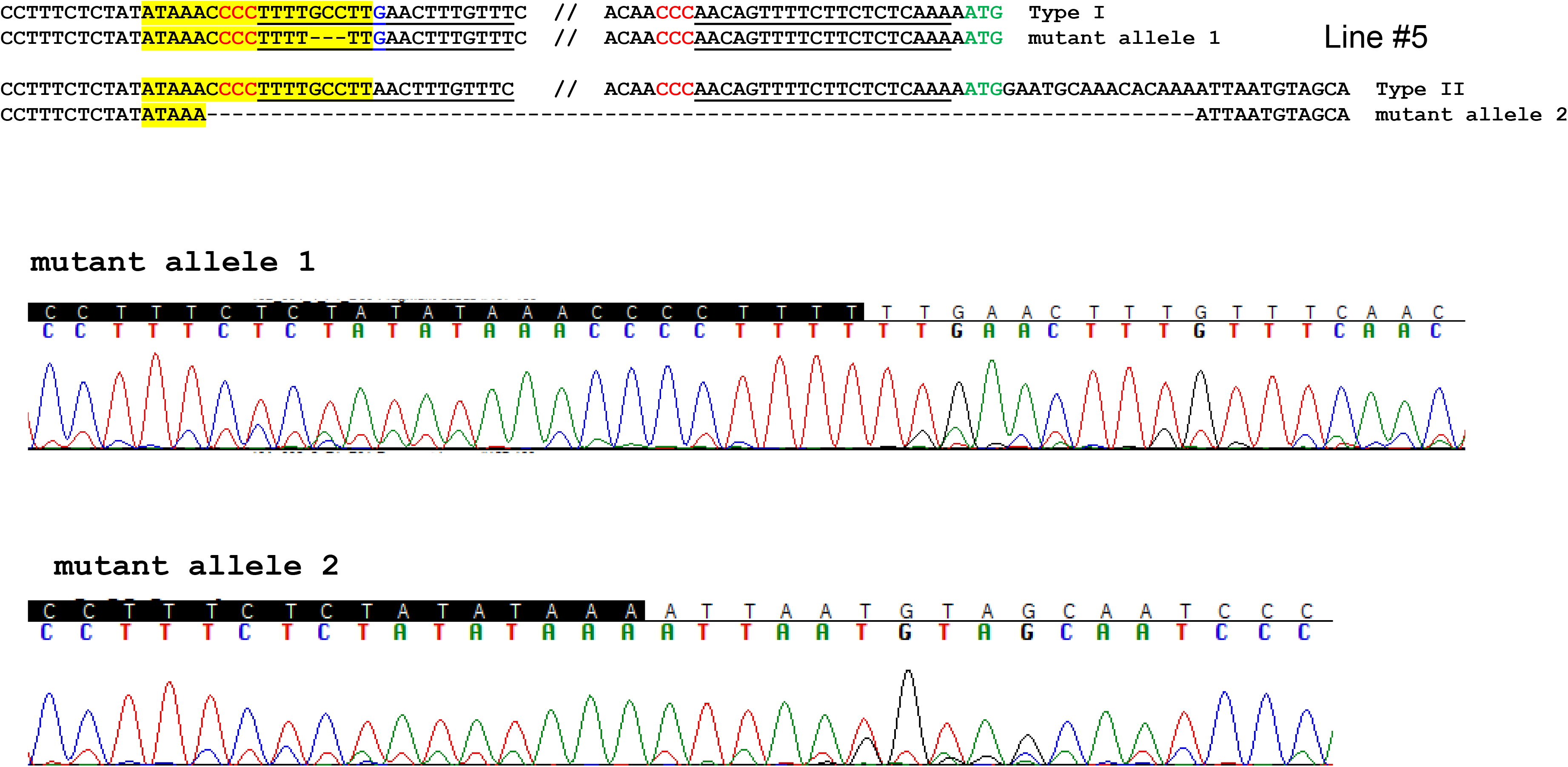
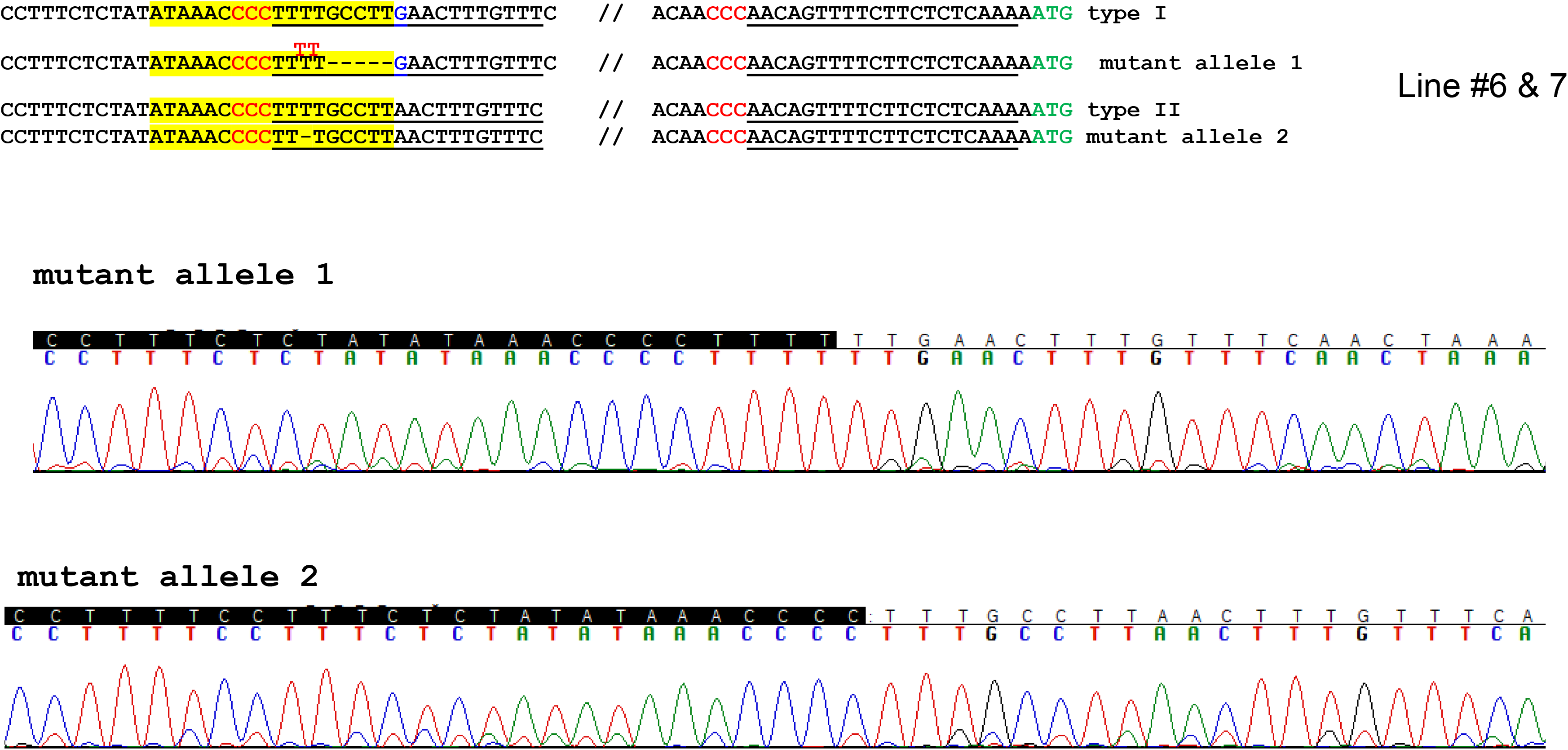
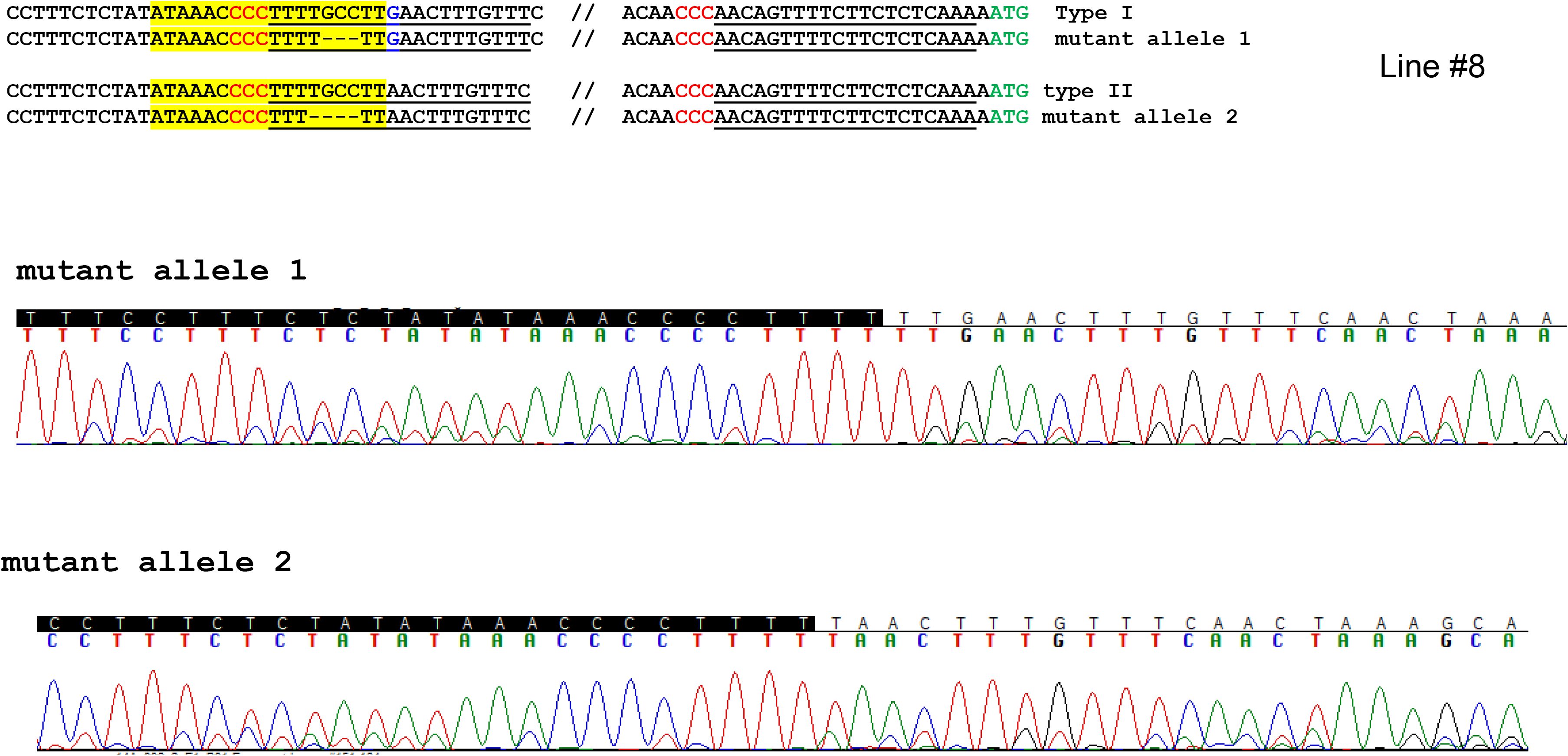
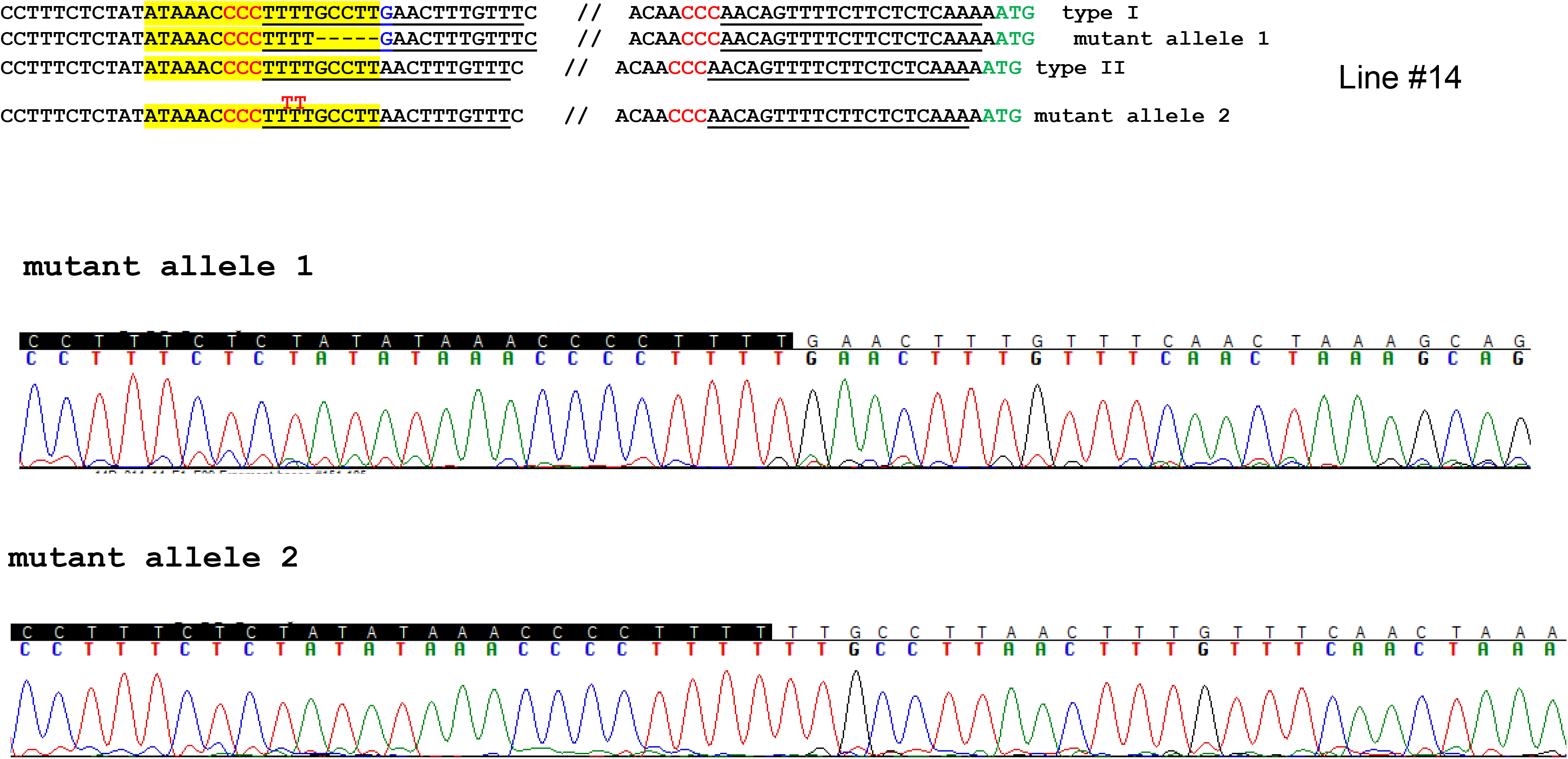
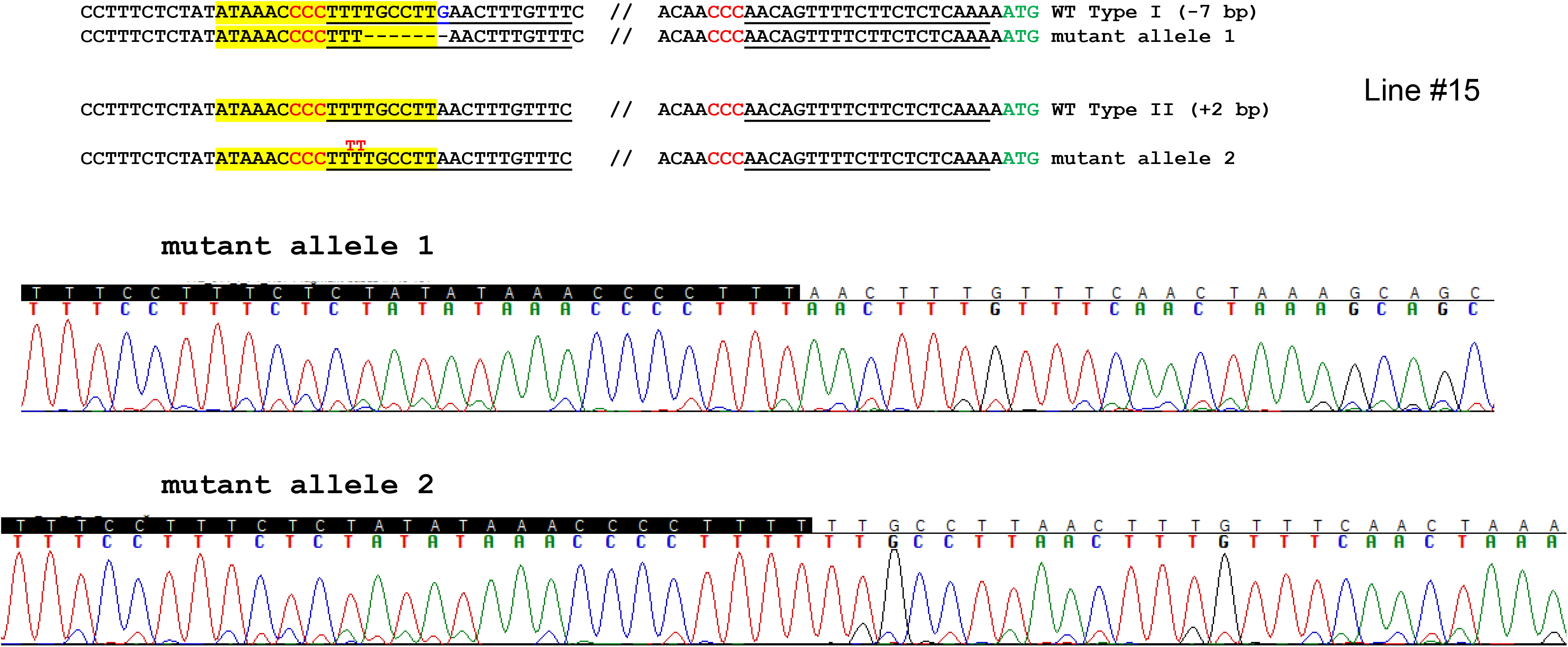
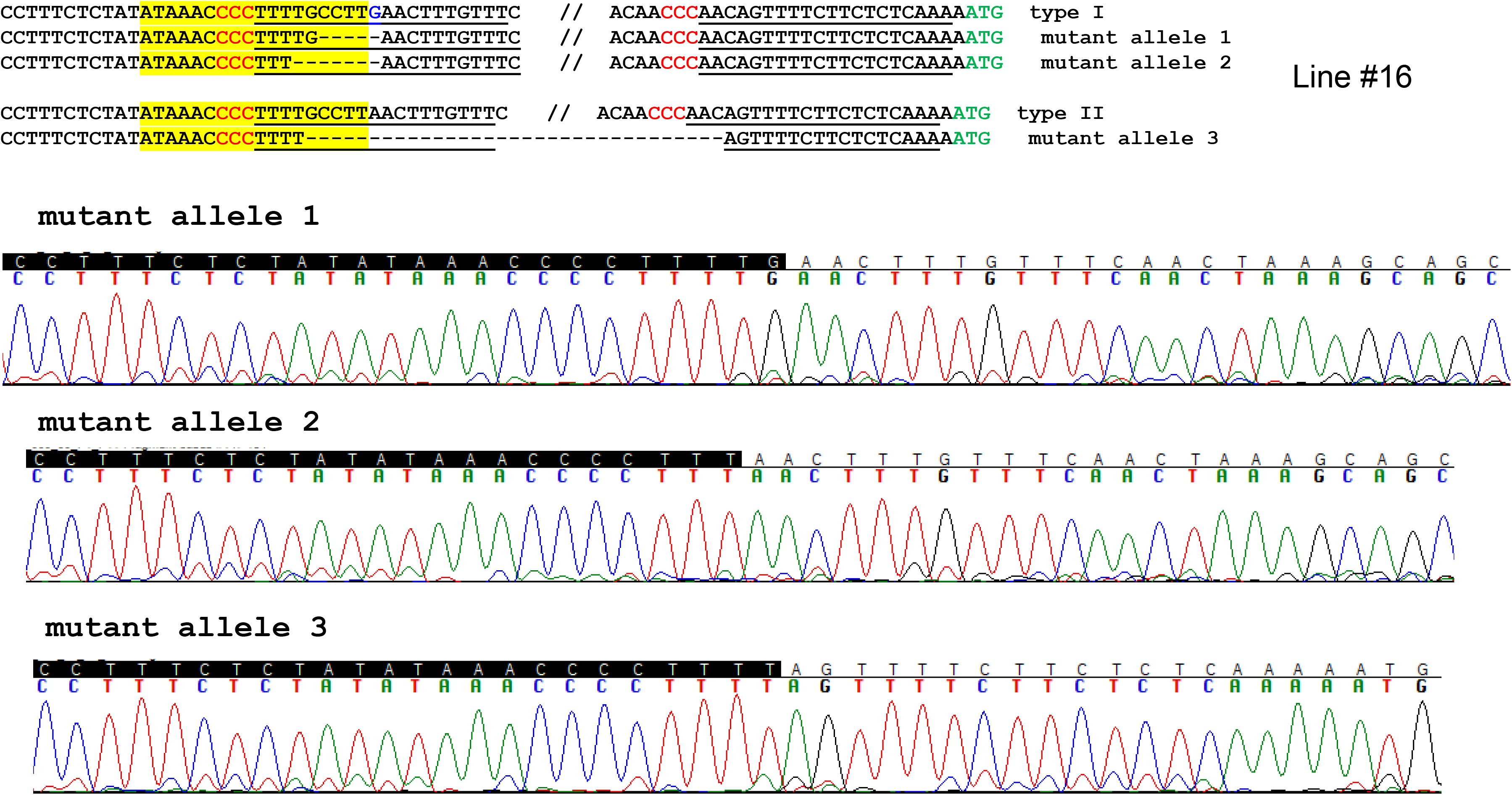
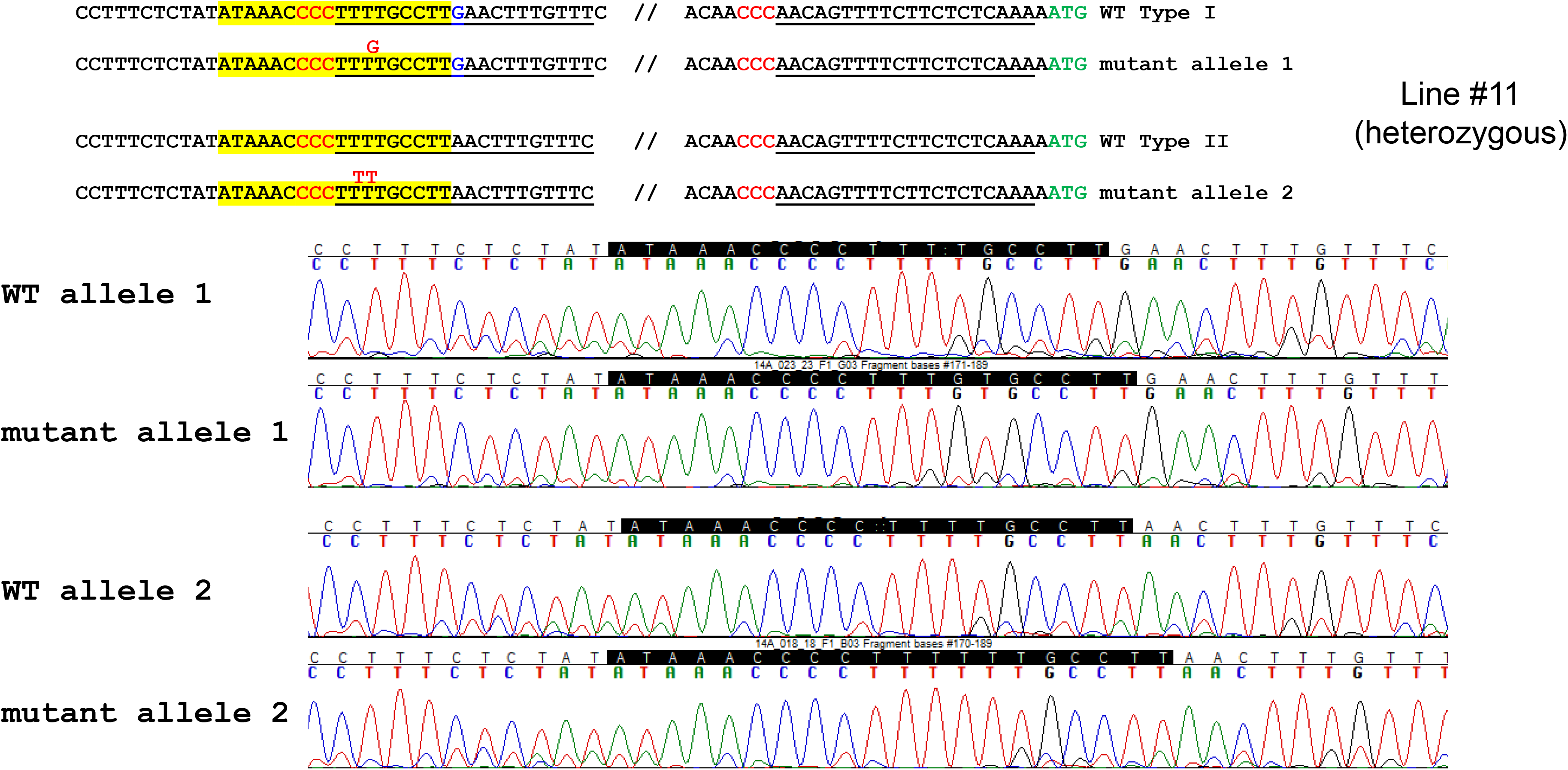
Sequence confirmation of the genome modified Hamlin sweet orange in the EBE region of CsLOB1.

---

**Supplementary Table 1.**
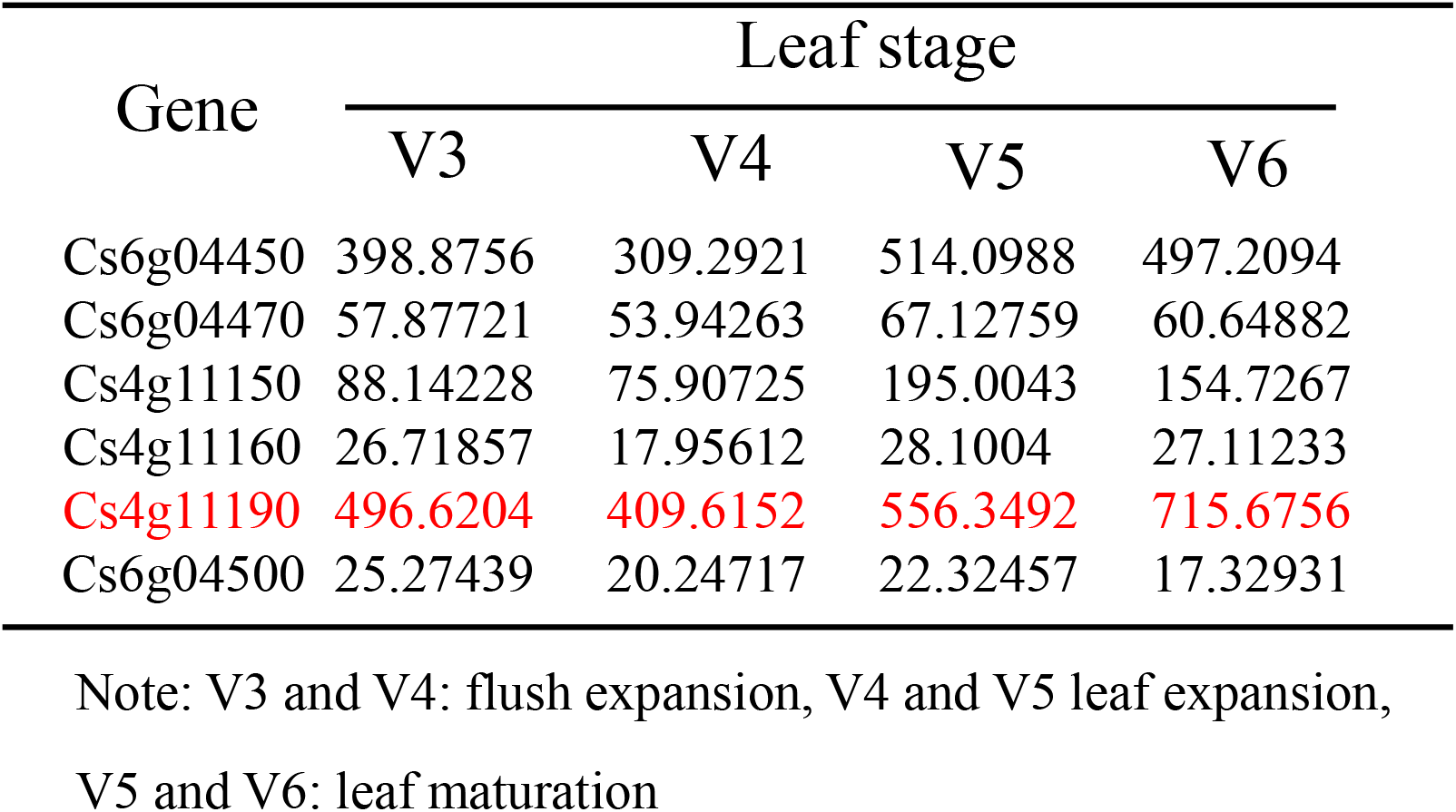
Expression levels of citrus ubiquitin genes at different leaf developmental stages from the RNA-seq data.

**Supplementary Table 2.**
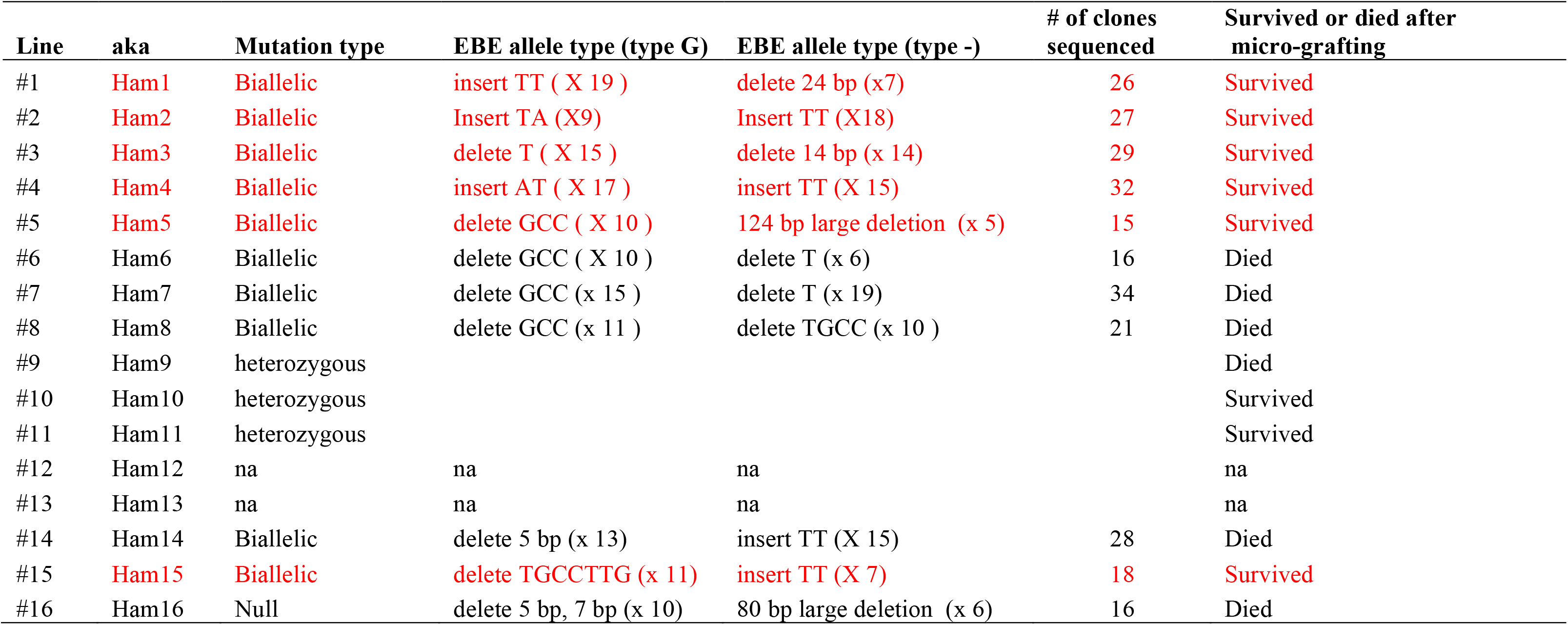
Genotyping of EBE mutants.

**Supplementary Table 3.**
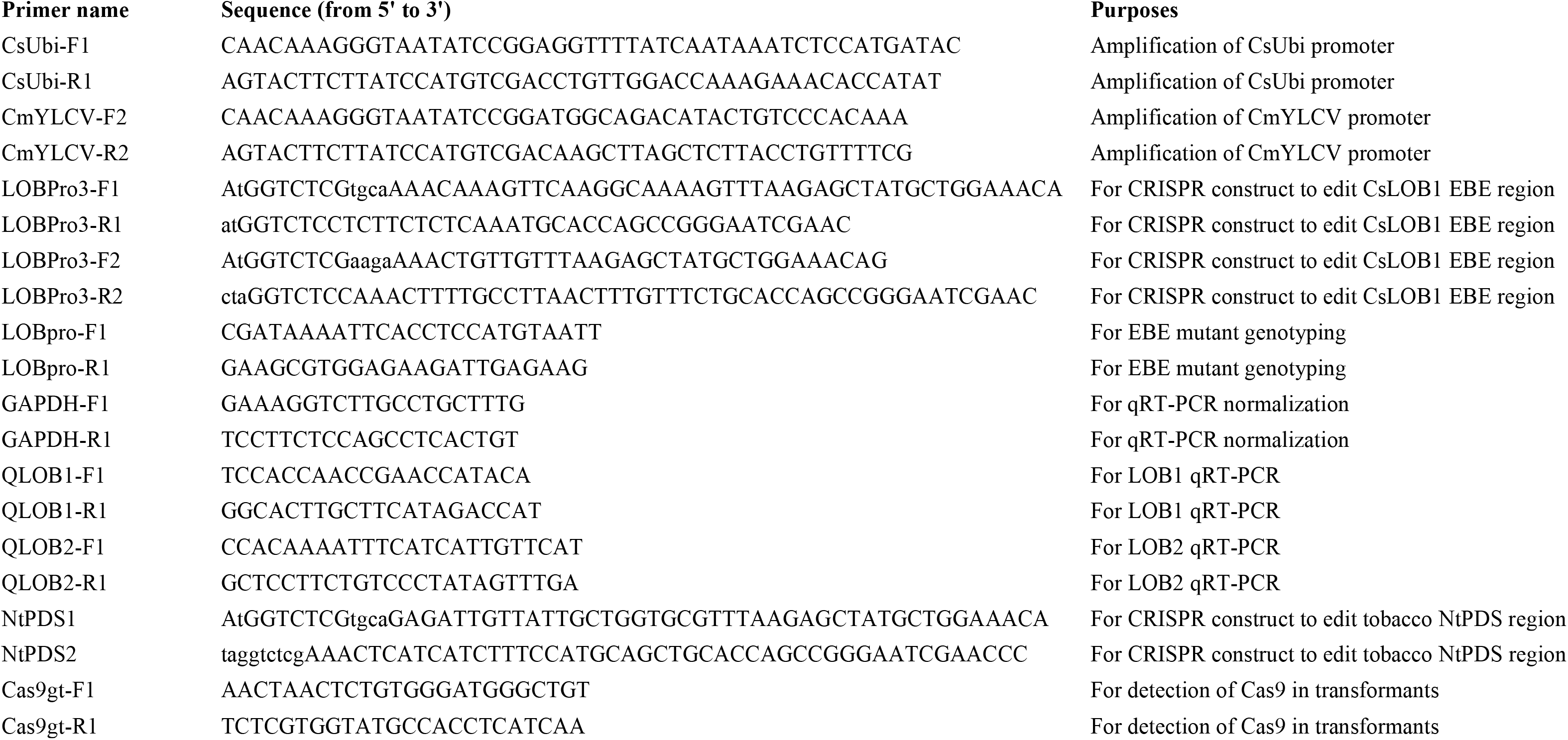
List of primers used in this study.

## Supplementary Data 1

### 35S enhancer-CmYLCV promoter

Yellow, 35S enhancer; **Black, CmYLCV Promoter**

GCATGCGGCGCGCCGATCGGCGCGCCAGATTTGCCTTTTCAATTTCAGAAAGAATGCTAACCCACAGATGGTTAGAGAGGCTTACGCAGCAGGTATCATCAAGACGATCTACCCGAGCAATAATCTCCAGGAAATCAAATACCTTCCCAAGAAGGTTAAAGATGCAGTCAAAAGATTCAGGACTAACTGCATCAAGAACACAGAGAAAGATATATTTCTCAAGATCAGAAGTACTATTCCAGTATGGACGATTCAAGGCTTGCTTCACAAACCAAGGCAAGTAATAGAGATTGGAGTCTCTAAAAAGGTAGTTCCCACTGAATCAAAGGCCATGGAGTCAAAGATTCAAATAGAGGACCTAACAGAACTCGCCGTAAAGACTGGCGAACAGTTCATACAGAGTCTCTTACGACTCAATGACAAGAAGAAAATCTTCGTCAACATGGTGGAGCACGACACACTTGTCTACTCCAAAAATATCAAAGATACAGTCTCAGAAGACCAAAGGGCAATTGAGACTTTTCAACAAAGGGTAATATCCGGATGGCAGACATACTGTCCCACAAATGAAGATGGAATCTGTAAAAGAAAACGCGTGAAATAATGCGTCTGACAAAGGTTAGGTCGGCTGCCTTTAATCAATACCAAAGTGGTCCCTACCACGATGGAAAAACTGTGCAGTCGGTTTGGCTTTTTCTGACGAACAAATAAGATTCGTGGCCGACAGGTGGGGGTCCACCATGTGAAGGCATCTTCAGACTCCAATAATGGAGCAATGACGTAAGGGCTTACGAAATAAGTAAGGGTAGTTTGGGAAATGTCCACTCACCCGTCAGTCTATAAATACTTAGCCCCTCCCTCATTGTTAAGGGAGCAAAATCTCAGAGAGATAGTCCTAGAGAGAGAAAGAGAGCAAGTAGCCTAGAAGTAGTCAAGGCGGCGAAGTATTCAGGCACGTGGCCAGGAAGAAGAAAAGCCAAGACGACGAAAACAGGTAAGAGCTAAGCTT

### 35S enhancer-CsUbi promoter

Yellow, 35S enhancer; **Black, CsUbi Promoter**

GCATGCGGCGCGCCGATCGGCGCGCCAGATTTGCCTTTTCAATTTCAGAAAGAATGCTAACCCACAGATGGTTAGAGAGGCTTACGCAGCAGGTATCATCAAGACGATCTACCCGAGCAATAATCTCCAGGAAATCAAATACCTTCCCAAGAAGGTTAAAGATGCAGTCAAAAGATTCAGGACTAACTGCATCAAGAACACAGAGAAAGATATATTTCTCAAGATCAGAAGTACTATTCCAGTATGGACGATTCAAGGCTTGCTTCACAAACCAAGGCAAGTAATAGAGATTGGAGTCTCTAAAAAGGTAGTTCCCACTGAATCAAAGGCCATGGAGTCAAAGATTCAAATAGAGGACCTAACAGAACTCGCCGTAAAGACTGGCGAACAGTTCATACAGAGTCTCTTACGACTCAATGACAAGAAGAAAATCTTCGTCAACATGGTGGAGCACGACACACTTGTCTACTCCAAAAATATCAAAGATACAGTCTCAGAAGACCAAAGGGCAATTGAGACTTTTCAACAAAGGGTAATATCCGGAGGTTTTATCAATAAATCTCCATGATACTTTTATTGAAAATTGTCATAAATTATCAACACGGATTATCAACCTAATTGAGAGTTTTCAATTTTATATTTTTTTAGTGATTGTAAAATTTCCCTCTTATTCCTAAATCACTATATTTATTTATGATTAATTCAATTACGCTACTAGAGTGATTCTTATAATTTTGGTTGAGTAGTTTTCACTTTCATCCAGCTCAAAAACTTAAAATTACAAATTGAAAAAAAAATTTGGATTTTATTATCAGAAATGCTGTTATAGGAATCTTTAAATTTTCGCAGATGTTCTTCTCAGAAAAACTTATTTAACATTTTTTTTAAGATAACTTTAAATCGTTAAAAATAATCCACAAATATACAAATAGAAAATTAGAATGATGAGCTACCGTGTGAGGTGTGGCACACTCGTATCGAAGCGCGTGACTTGAATGCTAAACAATATCCTAAAATCTGAAAGATATCCGCCTCTTCCGATCAGCCAACGGTTTTGCGAAACTACTATTGGCGTGTGGGCCAGGAAAGGGACACCGAATGTTAAAGACGTGGCATCAATGTGGTGGATGGAATTTGGGCAATCTCGTCATTTTAATTGTCAATCACGACTATAAAATGGAGGACCTCGAACCTCGGCTCCCCACCGTTTCTCAGATTATCTTCACCTTTAATTCGATAAAAGGCCTCCTTCTGTTCTCTCTGTCAAGGTAATAATGATTATAATTCATCTCTTTAGCATTTTGGTTTTTTTTTAAAATTAATATTTATTTTACCGATTATTCTAGCTGCTATTATGTCGTGCTTTGTCGATTAATTCGTAGATTTTTTTTTTTAGTTATTATTGTTTATCTTGATATGGTGATATGCTATTTATAAGCTTTTCATTAGCTGTAAACTTTTGTAGGCGATCGCTTATTTGTTGTTTGAGGTTATAATTATGGCGATTTAACACTCAAATCAGCCTCGTTGATTCTATCTAGGGTTTTGATTTAATGGTGTTATAGTGCTCTTTCGGAGATAATTTTGAATTTTTTGGTGTTAATTTTAGGCGTATCTTGTAATTCTATGAGACTTGTGAAAAAAAAATAATTAACAAAGAAAAAGGTATGCCGAAATCTTTACTTATGATTGATCGTTCTTGAAAATCTCTTTTCTATTTTTTGATCTATCTAGAGATCTACAACTTTTCTAAATTTTCTGTTAGTTTTTCCTTGAGAGACCTGCCCGAAAGTGTCTTTGAATTATGTAGTGACTACGTCTAACACAACTCTGTGTCGTTGGTTATGTTAATATTGTTTGTTTGGTTTGATTGTTTTGATTGTTTTGTATGGATTTATTCTGATGTGGTGTTTCTTTGGTCCAACAGGTCGAC

### CsU6 promoter (also known as CsU6-2 promoter)

CGCTCAGGAGCCGGTTGAATTTGATTGTTGTTTGATGTTTAGGTATGCTTACAATTTTTTACTAAATAAGGTAATAATTGTTCCTTTTATTTTATTCATTTAGCTTGGTACTTGTCGTAATCTATCTCCGTCAATGTCAGCTTCTTCTGTGTCGGAAGAAGCTGAAAAATGTTGTTGAAGGCCCTTTCATTTATTATTATCATTATTATGTTTAATTTTGTTCGGCAAATGTGTACTAATTTGGTTCGAATCGTTCGATCTCATAATACAAACATGTAATGAGTTATTTATGACGTCATGGATGCGGTTAGTTTGCGTGGTCTGAAATTTCAACCAAATTATTCAATCATGGTGGTGGCCGGCTGGTGCCTATCCCTTGATTGAAATATGCAATTAAAATTCTCGTAATAATGTTGATTTGTTCTATATACTTGAATTGATGGTATTAATTATTGTTATAAAAGCACTAACTTGTTTGAGAAAGGGGATAACTAAAAGGTAATAATAATAATAATAAAATAAAGAAAAGGCCCAACACATGGGCGCCCTATCTGTTGGGCCAAATGCCATAAGAGGATCCAGCTAGCCCGTTGAAGAAAATCCCACATCGAAAGAAAAAACTGAATAAACATGGTCTATATATACAAGGACTCCCAGGTTGGTTG

### CsU6 3’ region

**3’ region regulatory sequence of citrus CsU6 gene (downstream of PolyT)**

CCCCTGTTTTTTCCTAATTAGATTTCTTTCGGAGCTGTTCGAAGAACATGTTTTTGCCGTCTGACTCGTTTGTTATTGCCGTTCTAGTTAGTTTCAGCTATTGGTTTTCGGTTTCTTTCTGTTTTGAAGGGTAATAGTTTGGAAAATATAATACAAATTAATTTAGGATTAAAAAGTAATAAAACAATATTATCTAGA

